# A general method for bootstrapping dense 3D segmentations from sparse 2D annotations

**DOI:** 10.1101/2024.06.14.599135

**Authors:** Vijay Venu Thiyagarajan, Arlo Sheridan, Kristen M. Harris, Uri Manor

## Abstract

Volumetric electron microscopy (vEM) provides unparalleled insight into cellular architecture, transforming our understanding of fundamental biological processes. Precise quantification of volumetric bioimaging data requires 3D instance segmentation, but the time and cost of manual ground-truth annotation present a major bottleneck for training accurate machine learning models. Our lightweight two-stage method generates complete 3D instance segmentations from sparse 2D annotations. A 2D network predicts dense boundaries on every section from minimal 2D labels. A 3D network pre-trained on synthetic data infers connectivity between sections. Across the imaging modalities and biological structures tested, our method generates base segmentations with minutes of human effort per volume. Models trained on these 2D→3D pseudo ground-truth segmentations approach the accuracy of fully supervised models while reducing human annotation time by four orders of magnitude. This method does not require dense 3D labeling and thus democratizes scalable volumetric analysis across imaging modalities in basic science and biomedical applications.

## Introduction

Volumetric electron microscopy (vEM) is generating increasingly large image datasets, and the three-dimensional (3D) information in these volumes is revolutionizing our understanding of basic biological principles and disease processes^1–4^. Historically, analyses of these datasets required tedious manual annotation of the objects of interest throughout entire tissue volumes. Usually, annotation involves an expert using a computer mouse or pen to painstakingly “paint” pixels of exact membrane boundaries in grayscale EM images visualized on a computer monitor. Deep learning-based segmentation models obviate manual annotation, but training high-accuracy models requires large amounts of ground-truth data that is expensive and time-consuming to generate^5^. Diverse ground truth from different imaging conditions, tissue types, and anatomical regions is required for models to generalize, further amplifying the costs. Segmentation foundation models such as DINOv3^6^, SAM^7^, microSAM^8^, and CellSAM^9^, and generalist models such as Cellpose^10^ produce 2D segmentations across many imaging domains. However, these models often perform poorly on rare or complex structures, particularly in vEM, necessitating model adaptation, additional annotation, and iterative proofreading^11,12^.

Several lines of work aim to reduce annotation effort for 3D segmentation. Strategies relying on consensus among multiple predictions, with or without pretrained models or orthogonal planar slicing, offer theoretical generalizability^10,13–16^. Other work has demonstrated pipelines that learn to correct incomplete annotations such as skeletons and seeds^17^, as well as weakly supervised methods^18^. These approaches have not yet been extended to 3D segmentation of complex structures such as the brain. Methods built for neurons in EM still train on dense 3D labels, even when a lightweight 2D network predicts the 3D affinities for faster inference^19^. Moreover, dense and diverse ground truth remains essential for rigorous model evaluation, even for self-supervised approaches.

An ideal method would produce dense 3D segmentations from sparse 2D annotations, as annotation is much easier in 2D than in 3D, and easier when sparse than when dense. Here, we present a lightweight two-stage 2D→3D segmentation pipeline that generates relatively accurate, dense 3D instance segmentations from sparse 2D annotations, providing high-quality pseudo ground truth suitable for model bootstrapping and refinement. During the first stage, a 2D network trained on sparsely painted 2D labels predicts the entirety of dense boundaries on every section of a 3D volume. In the second stage, a 3D network pre-trained on synthetic geometric shapes then infers inter-section connectivity across the stacked 2D predictions from the first stage. Importantly, the model does not require absolute pixel-perfect input annotations, greatly reducing the effort and expertise required to generate large amounts of pseudo ground-truth data.

We validated this framework across 11 diverse segmentation tasks on public datasets, encompassing multiple imaging modalities and biological structures including whole cells, neuronal processes, organelles, and synapses. Downstream 3D models trained on our pseudo ground truth achieved performance comparable to fully supervised models trained on expert manual annotations, while reducing human annotation time by four orders of magnitude. We provide the complete framework as an open-source command-line tool and an interactive plugin for the napari graphical user interface^20^ to ensure accessibility.

## Results

### Dense 3D segmentation from sparse 2D annotation

The 2D→3D method converts sparse 2D labels into a dense 3D instance segmentation in two steps (Fig. 1a). First, a 2D U-Net trained on a few sparsely labeled sections predicts dense boundaries on every section. Then a lightweight 3D U-Net^21^ trained only on synthetic 3D boundaries infers how those predictions connect across sections. Thus, the method obviates the need for manual 3D annotation of boundaries.

**Fig. 1.**
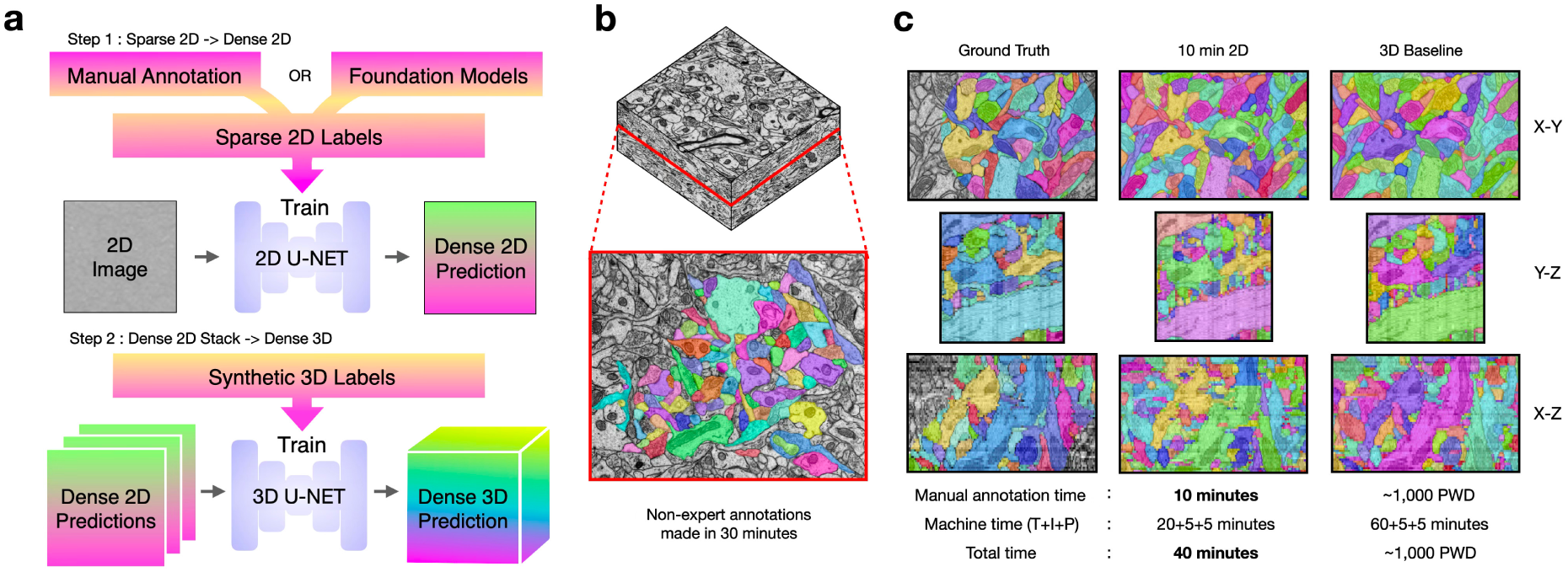
The 2D→3D method generates dense 3D segmentations from sparse 2D annotations. **a**, Overview of the method. Step 1 trains a 2D U-Net on sparse 2D labels, from manual annotation or from a segmentation foundation model, to predict dense 2D boundaries on every section. Step 2 trains a 3D U-Net on synthetic 3D labels to infer connectivity through a stack of 2D predictions and output 3D predictions. **b**, Sparse non-expert annotations created in 30 minutes on one section of HARRIS-15. **c**, Orthogonal views of HARRIS-15 comparing the ground truth, the 2D→3D result from 10 minutes of sparse 2D annotation on one section and the 3D baseline result from the full dense 3D annotation. The dense 3D labels trained the baseline and scored both results, while the 2D→3D method saw only 10 minutes of the 2D labels in b. Total time to segmentation is the sum of human annotation time and machine time. Machine time is the sum of training (T), inference (I) and post-processing (P) times. The dense annotation time of the baseline is given in person-workdays (PWD; 5 h per workday).

Ten minutes of non-expert painting on a single section of HARRIS-15 hippocampal neuropil^22^ produced a dense 3D segmentation comparable to that of a 3D baseline model trained on the full expert ground truth (Fig. 1b, c). The total time for segmentation, including training, inference and post-processing, was 40 minutes. Dense annotation for the baseline took about 1,000 person-workdays (Fig. 1c). HARRIS-15 is the most challenging of our datasets because of its low axial resolution and the diversity of intertwined axonal, dendritic, and glial processes.

We obtained comparable results across six public datasets spanning serial-section transmission EM, focused ion beam SEM and fluorescence microscopy: HARRIS-15, FIB-25^23^, CREMI-A, CREMI-B and CREMI-C^24^, and EPI^25^ (Supplementary Table 1). For each dataset, the 2D→3D method was trained on sparse labels from Volume 1 and evaluated on Volume 2. Simulated sparse training labels contained 1 to 100 labeled objects, with or without disk masks that mimic brush strokes (Supplementary Fig. 1; Supplementary Note 1). We also used training labels from 10 or 30 minutes of non-expert painting (Supplementary Fig. 2). At every annotation level, normalized variation of information (VOI)^26,27^ approached that of the 3D baseline trained on dense ground truth. Accuracy improved as more objects or sections were labeled (Fig. 2; Supplementary Figs. 3–5). Labels that sampled the diversity of structures in a volume outperformed labels concentrated on a few structures.

**Fig. 2.**
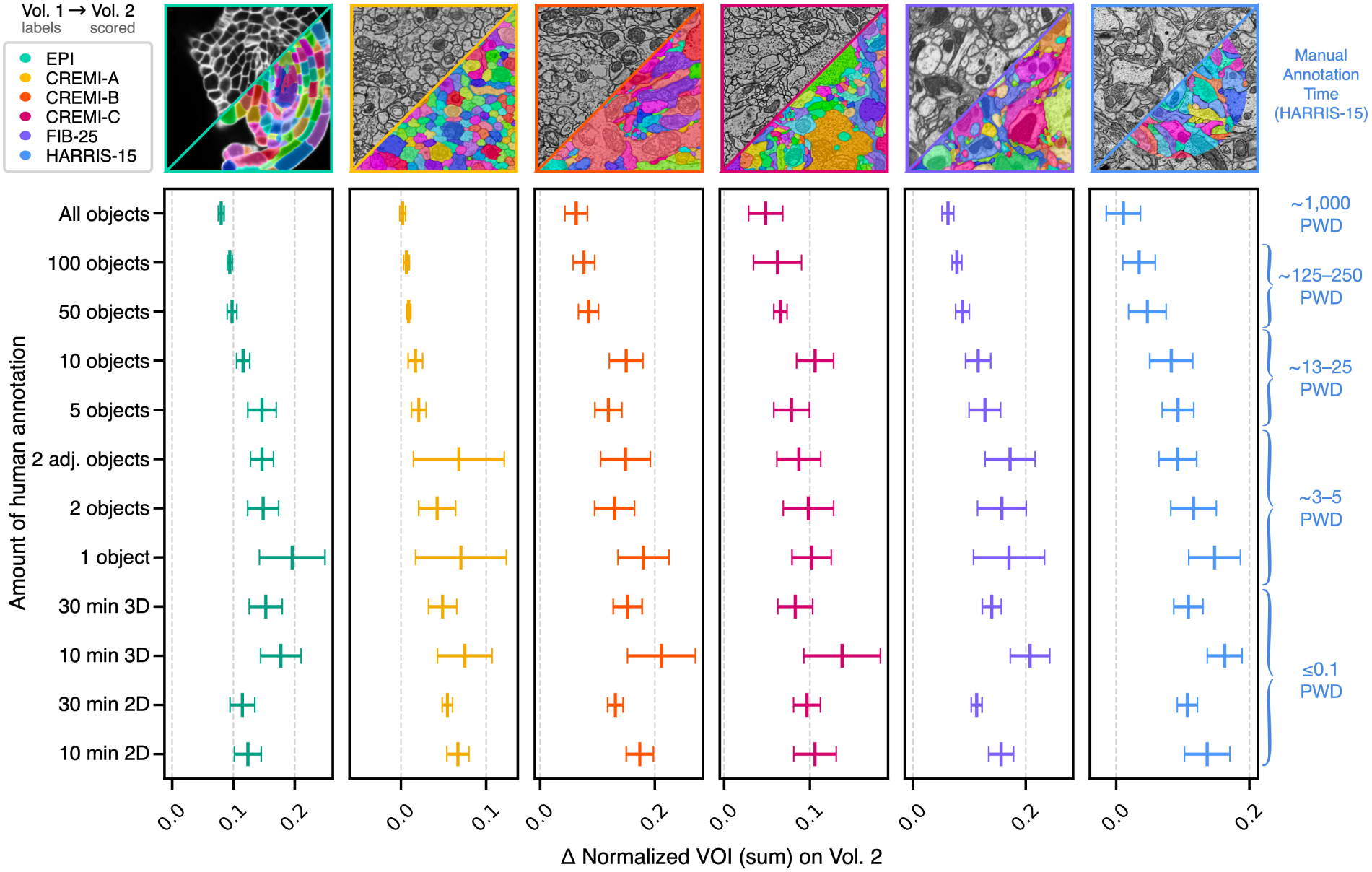
Accuracy of the 2D→3D method at each annotation level on six datasets. Deviation (Δ) of the normalized variation of information (VOI) sum from the 3D baseline for EPI, CREMI-A, CREMI-B, CREMI-C, FIB-25 and HARRIS-15 (left to right). Rows are the annotation levels used to train the 2D network: all objects; 100, 50, 10, 5, 2 adjacent, 2 or 1 labeled objects; and 30 or 10 minutes of non-expert painting in 3D or 2D. Lower is better. Δ is the VOI sum of the 2D→3D method minus that of the 3D multitask local shape descriptor (MTLSD) model trained on the dense ground truth of the same repetition. Points are means and bars are ±1 standard deviation (s.d.) over n = 12 differences (4 types of 2D prediction × 3 repetitions). The 4 types are 2D affinities, 2D LSDs, and the affinities and LSDs of the 2D MTLSD network (Methods). Labels on the right give the approximate human annotation time of each level for HARRIS-15 in person-workdays (PWD; 5 h per workday; Methods).

### Synthetic training of the 3D network

Two training choices, masked loss and synthetic data, remove the need for 3D annotation (Fig. 3a). The 2D U-Net^28^ is trained with a loss restricted to labeled pixels, so unlabeled regions are neither penalized nor learned as background. Even so, the network predicts complete boundaries on every section, encoded as 2D local shape descriptors (LSDs)^29^ (Fig. 3b). The 3D U-Net uses entirely synthetic training data, with input (stacked 2D LSDs) and target (3D affinities) computed from synthetic 3D label volumes (Fig. 3c). These volumes are generated by growing random seeds or by applying a watershed to smoothed noise (Fig. 3d; Methods). The 3D networks of the 2D→3D method learn geometric priors for connectivity between sections from synthetic shapes alone and are never trained on the target data.

**Fig. 3.**
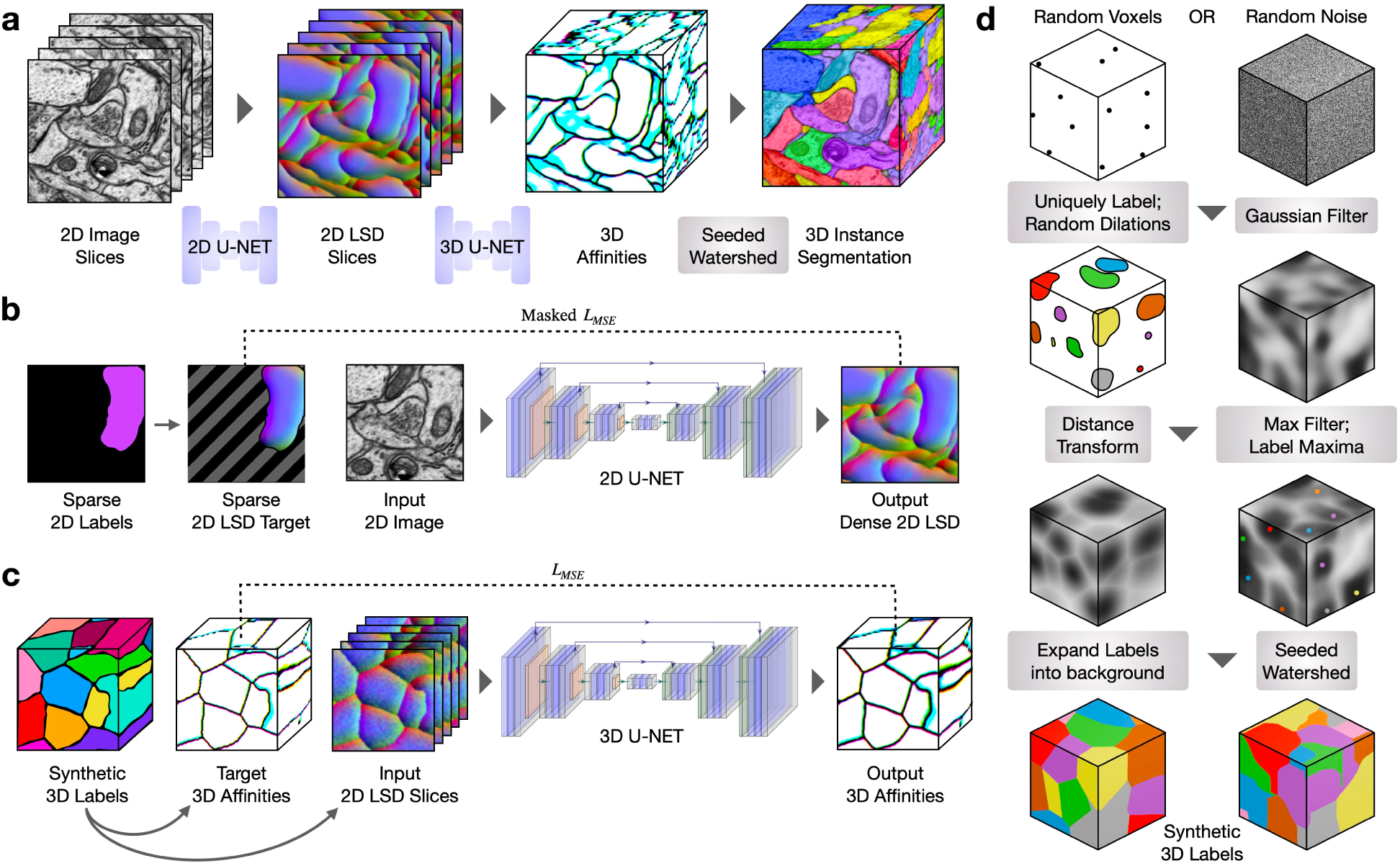
The 2D→3D pipeline. **a**, Inference. Five example sections are shown for simplicity. The trained 2D U-Net predicts 2D local shape descriptors (LSDs) from sections of a 3D image volume, input one at a time. The trained 3D U-Net infers 3D affinities from the stack of 2D LSDs. LSDs and affinities are RGB encoded for visualization. Seeded watershed and hierarchical agglomeration of the affinities generates the 3D segmentation. **b**, Sparse 2D to dense 2D training. 2D images with sparse labels train a 2D U-Net to predict dense 2D LSDs. Background regions of the target 2D LSDs, marked by diagonal gray stripes, are masked out of the loss. All networks are trained with a weighted mean-squared-error loss between predictions and targets, denoted L_MSE_. **c**, Stacked dense 2D to dense 3D training. Target 3D affinities and input 2D LSD slices are simulated from synthetic 3D labels. **d**, Generation of synthetic 3D labels by a series of morphological operations and transformations applied to a random array of noise or foreground voxels.

In the experiments of Fig. 2, seeded watershed and hierarchical agglomeration of the predicted 3D affinities gave the final instances. The comparison and generalization experiments below used mutex watershed (Methods). Consensus methods instead fuse pre-segmented 2D sections^13,14^. Here the 3D step is learned, runs blockwise on large volumes and needs only standard post-processing. The 3D network has 0.26 million parameters, and every network in the pipeline trains in less than 3 GB of GPU memory (Supplementary Table 2; Supplementary Notes 2 and 3).

### Comparison with Cellpose and uSegment3D

With 5 minutes of annotation, the 2D→3D method approached the accuracy of an established pipeline that relies on far more labeled data (Fig. 4). On an EPI test volume (Methods), we compared it with Cellpose^10^ run on all three orthogonal views of the volume. uSegment3D^13^ then fused the 2D predictions into a 3D consensus segmentation.

**Fig. 4.**
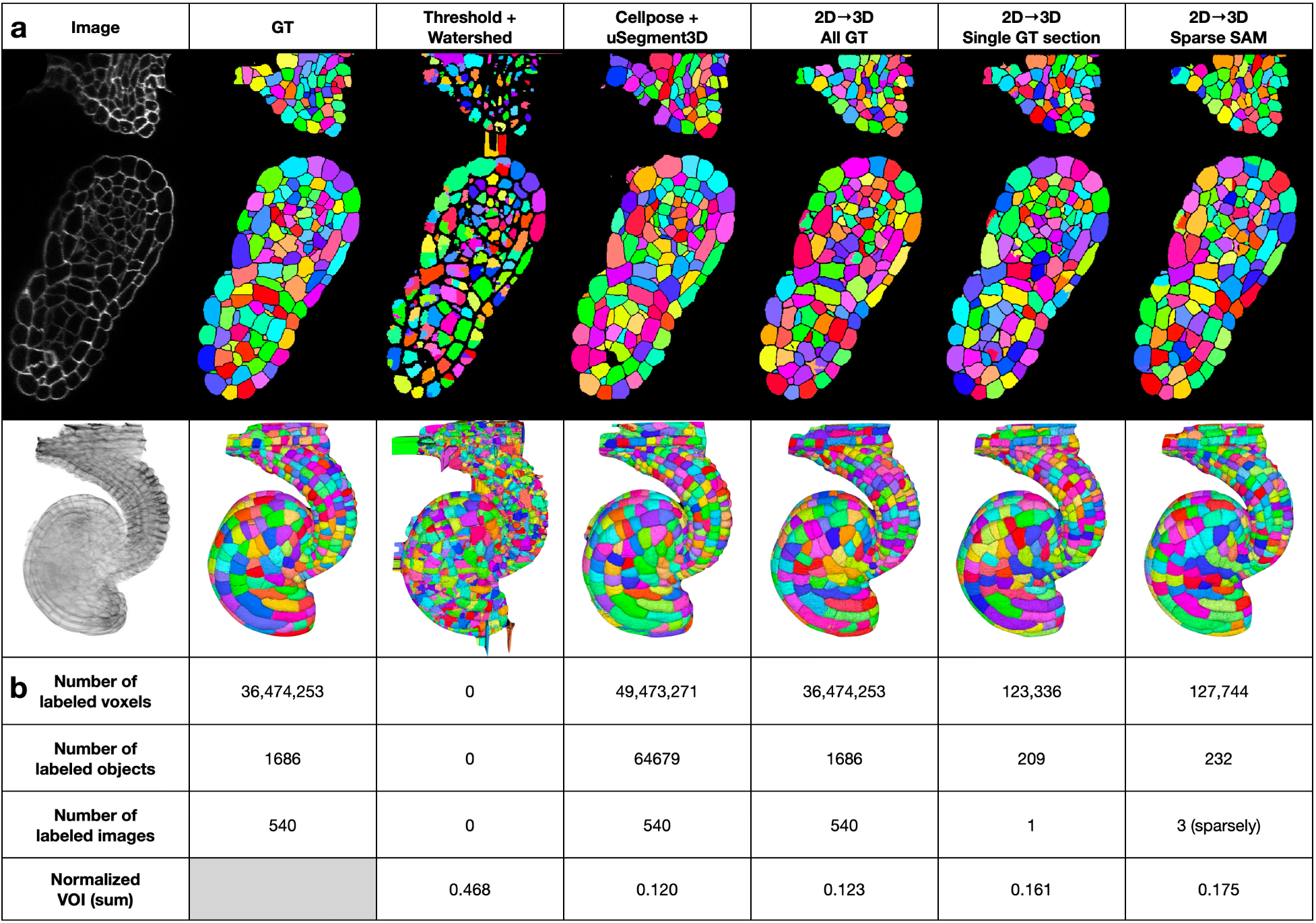
Comparison of the 2D→3D method with Cellpose + uSegment3D on an EPI volume. **a**, Representative x-y sections (top) and 3D mesh renderings (bottom) of the EPI test volume: the ground-truth (GT) labels and the segmentations of the annotation-free Otsu threshold and watershed baseline (Threshold + Watershed), of Cellpose + uSegment3D and of the 2D→3D method at each level of supervision (all GT, one GT section, or sparse Segment Anything Model (SAM) labels on three sections). **b**, Quantitative comparison. For each method: the number of labeled voxels, labeled objects and labeled images, and the normalized variation of information (VOI) sum against the ground truth. The labeled voxels, objects and images are the training labels of each method. For Cellpose + uSegment3D, they are the training set of the Cellpose model (Methods). Lower is better. One segmentation per method; no error bars.

Normalized VOI (sum) was 0.120 for Cellpose + uSegment3D and 0.123 for the 2D→3D method trained on the dense ground truth of the same volume. Trained on a single labeled section, the 2D→3D method scored 0.161. Trained on sparse SAM^7^ labels drawn on three sections in 5 minutes, it scored 0.175, using about 300-fold fewer labeled voxels than the dense ground truth. An annotation-free Otsu threshold and watershed baseline scored 0.468 (Fig. 4a, b).

### Generalization across imaging modalities and targets

We also tested the same pipeline qualitatively on five further public datasets (Fig. 5), with four target types: neurons, synaptic clefts, mitochondria, and whole cells. These datasets span expansion light microscopy, live-cell fluorescence time-lapse imaging and electron microscopy. Each segmentation needed only 10 minutes of SAM-assisted labeling or two to ten sections of the published ground truth, and the results were visually consistent with the published proofread or manual segmentations. Neurons were segmented in expansion-microscopy LICONN^30^ volumes of mouse hippocampal CA1 (10 minutes, SAM-assisted; Fig. 5a). Synaptic clefts were segmented in serial-section TEM of *Drosophila* neuropil (CREMI-C^24^; two sections of the published ground truth; Fig. 5b). Mitochondria were segmented in multi-beam SEM of human cortex (MitoEM-H^31^; two sections; Fig. 5c). Neurons were also segmented in 18-channel expansion light microscopy of mouse hippocampus (PRISM^32^; ten sections; Fig. 5d). Huh7 cells in live-cell fluorescence time-lapse imaging were segmented as 2D+t (Fluo-C2DL-Huh7^33^; two frames; Fig. 5e). Imaging parameters for all datasets are listed in Supplementary Table 1.

**Fig. 5.**
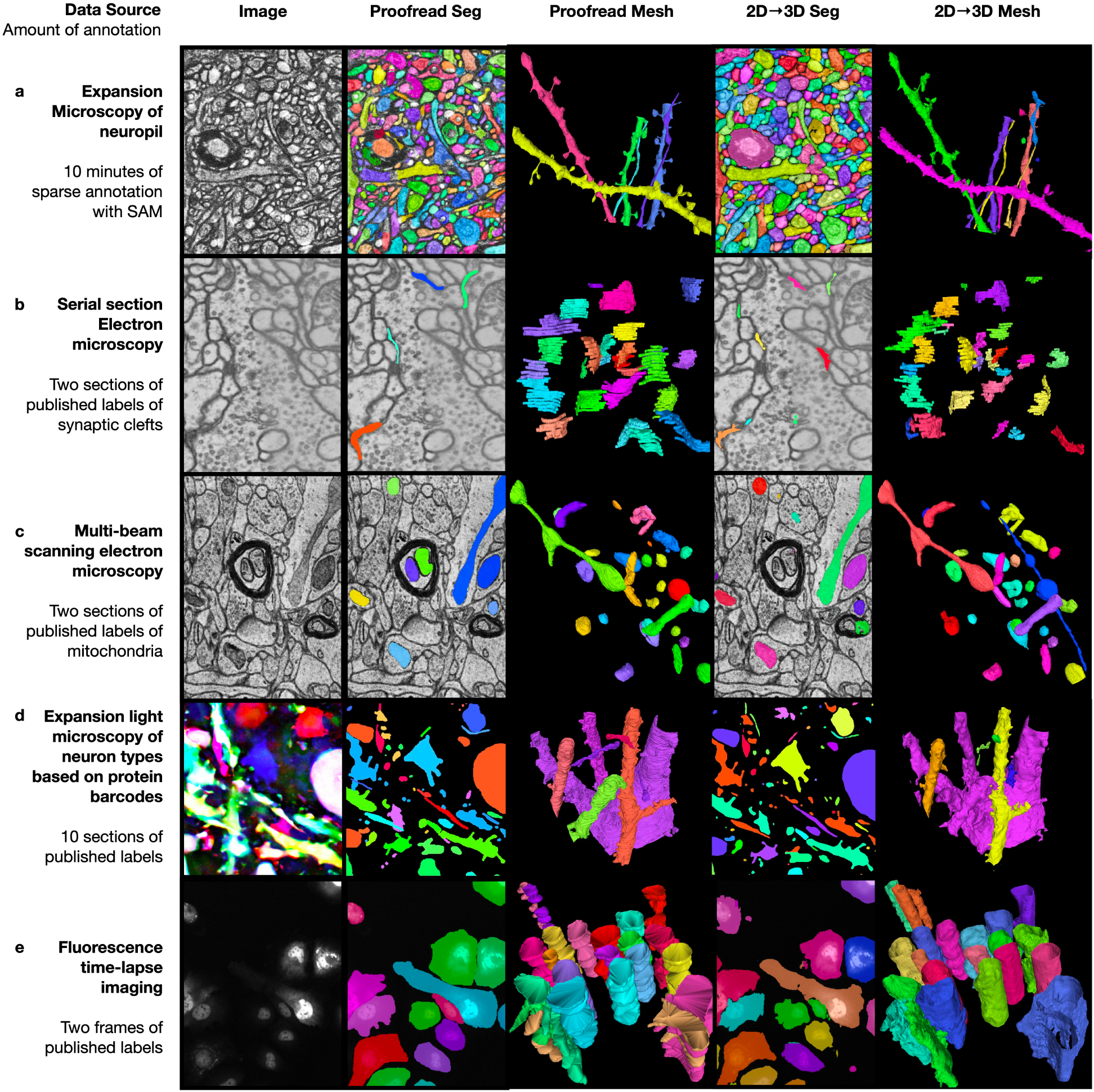
The 2D→3D method generalizes across imaging modalities and segmentation targets. Each row shows one dataset: a representative 2D image (column 1), the published proofread, manual or ground-truth segmentation (column 2), the published 3D mesh (column 3), the 2D→3D segmentation (column 4) and the 2D→3D mesh (column 5). **a,** Neurons in an expansion-microscopy LICONN^30^ volume of mouse hippocampal CA1 based on 10 minutes of SAM-assisted annotation. **b,** Synaptic clefts in serial-section transmission EM of *Drosophila* neuropil (CREMI-C^24^), from two sections of the published ground truth. **c,** Mitochondria in multi-beam scanning EM of human cortex (MitoEM-H^31^), from two sections of the published ground truth. **d,** Neurons in 18-channel expansion light microscopy of mouse hippocampus (PRISM^32^), from ten sections of the published ground truth. **e,** Huh7 cells in live-cell fluorescence time-lapse imaging (Fluo-C2DL-Huh7^33^), segmented as 2D+t, from two frames of the published ground truth. SAM, Segment Anything Model; EM, electron microscopy.

### Bootstrapping 3D models from pseudo ground truth

Unproofread 2D→3D segmentations were effective training data for a conventional 3D segmentation model (Figs. 6 and 7). Because the 2D→3D output is a dense 3D segmentation, it can replace manual ground truth for any 3D model that trains on dense labels. We tested this with a 3D multitask LSD (MTLSD) network^29^ reimplemented from the CREMI leaderboard. We left out dataset-specific optimizations so that the pipeline was identical across the six datasets.

**Fig. 6.**
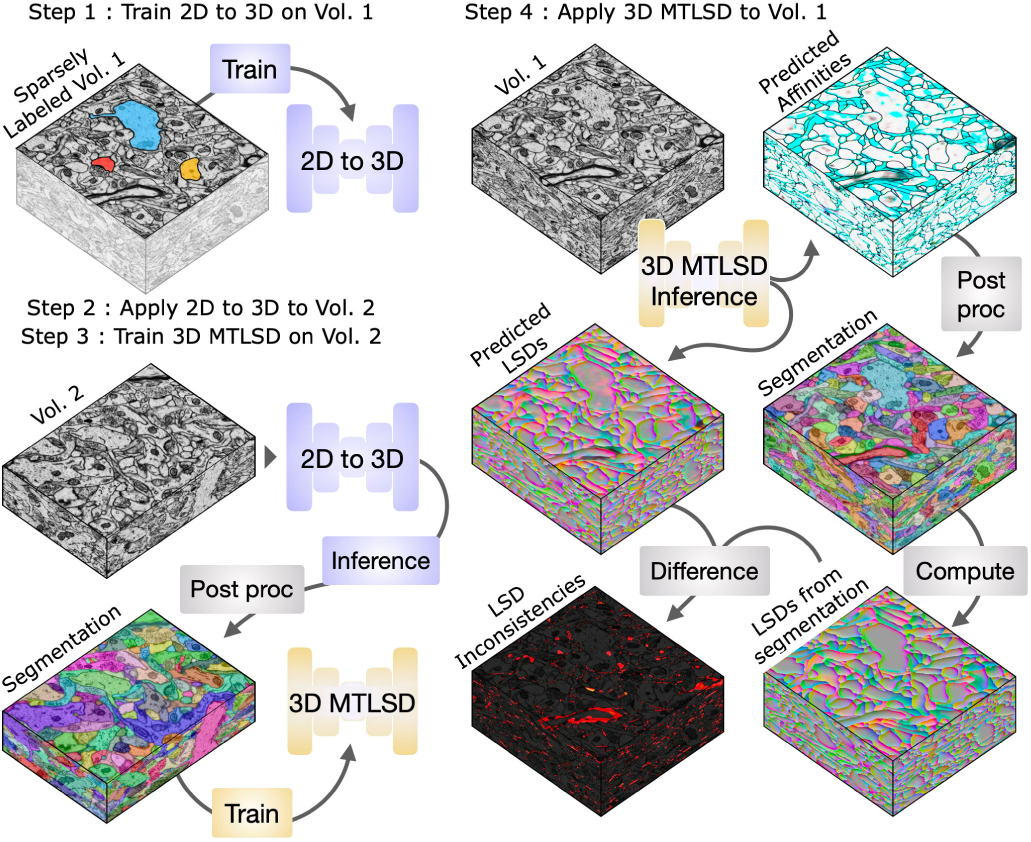
Bootstrapping 3D models from 2D→3D pseudo ground truth. Sparse 2D labels on Volume 1 train the 2D→3D method, which segments Volume 2. This unproofread segmentation is the pseudo ground truth that trains a 3D network to predict affinities and local shape descriptors (LSDs) together (multitask LSD, MTLSD). The trained MTLSD network then predicts 3D affinities and LSDs on Volume 1. LSD inconsistencies are the element-wise difference between the predicted LSDs and the LSDs computed from the final segmentation (Supplementary Fig. 9).

**Fig. 7.**
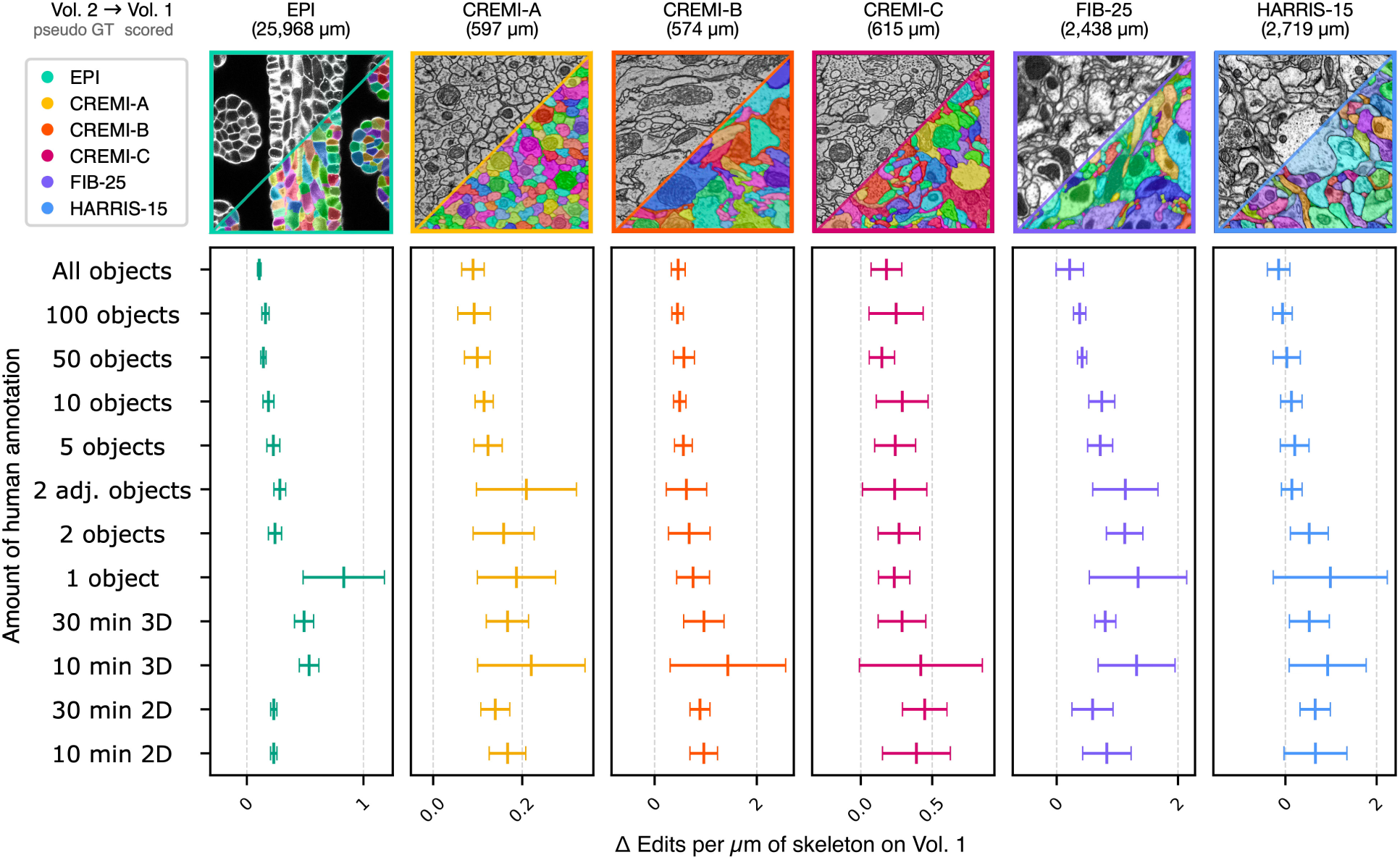
Accuracy of bootstrapped 3D models trained on the 2D→3D pseudo ground truth. Deviation (Δ) of the number of split and merge edits needed per micron of skeleton path length (µm^−1^) from the non-bootstrapped 3D multitask local shape descriptor (MTLSD) network trained on dense ground truth, for six datasets (total ground-truth path length in parentheses) and each annotation level of the pseudo ground truth. Lower is better. Points are means and bars are ±1 s.d. over n = 12 bootstrapped models (4 pseudo ground-truth network variants × 3 repetitions). The 4 variants differ in the type of 2D prediction (Fig. 2) that generated the pseudo ground truth. All means lie above zero except for HARRIS-15 with 100 or more objects, where the bootstrapped model outperformed the baseline. The manually annotated ground truth covers only a cylindrical subregion of Volume 2, whereas the pseudo ground truth segments all of Volume 2 and so provides more training labels.

The three-step workflow used sparse labels on Volume 1 to train the 2D→3D method, which segmented Volume 2 (Fig. 6; Supplementary Note 4). We used this segmentation, without proofreading or refining, to train the 3D MTLSD network. The trained network then segmented Volume 1 for evaluation against its dense ground truth. We also tested a 3D model bootstrapped through an intermediate subvolume. Training the first stage of the model on the evaluation subvolume itself gave no systematic advantage over training on an independent subvolume (Supplementary Fig. 6; Supplementary Note 5).

Across all datasets and annotation levels, bootstrapped models approached the accuracy of 3D models trained on dense expert ground truth (Fig. 7; Supplementary Figs. 7 and 8). Accuracy was measured as the number of split and merge edits needed per micron of skeleton (min-cut metric, MCM^29^). In HARRIS-15, pseudo ground truth from 10 minutes of non-expert annotation on one section required about 2.6 edits per µm, compared with about 1.8 for pseudo ground truth from all objects (Table 1). Regions where the predicted LSDs disagree with LSDs recomputed from the segmentation (LSD inconsistencies) mark where residual errors lie (Supplementary Fig. 9), conveniently highlighting targets for further proofreading and bootstrapping.

**Table 1.** Total reconstruction time for bootstrapping HARRIS-15. Edit counts and time estimates for 3D multitask local shape descriptor (MTLSD) models bootstrapped from 2D→3D pseudo ground truth at three annotation levels: all objects of Volume 1, 10 objects, or 10 minutes of non-expert 2D painting. Edits are the split and merge corrections needed to match the ground-truth skeletons of Volume 1, assessed using the min-cut metric (MCM). **Time assumptions.** PWD, person-workdays; 1 PWD = 5 h of annotation. False-split correction takes 0.1 min per error in every scenario; false-merge correction takes 1, 3, or 10 min per error, as indicated. Total reconstruction time = manual annotation + machine computation + proofreading. The dense annotation estimate of 1,000 PWD is described in Methods. **Edit-count reporting.** Counts are means across 12 models. Each metric was selected at its best checkpoint, so total edits need not equal the sum of the separately reported split and merge corrections.

| Measure |  | All objects | 10 objects | 10 minutes 2D painting |
|---|---|---|---|---|
| **A Corrections required** |  |  |  |  |
| Merge edits to correct false splits |  | 3,052 | 3,589 | 4,850 |
| Split edits to correct false merges |  | 1,822 | 2,027 | 2,159 |
| **Total edits** |  | **4,964** | **5,724** | **7,157** |
| **B Annotation and computation time in PWD** |  |  |  |  |
| Manual annotation |  | 1,000 | 25 | **0.03** |
| Machine computation |  | 0.3 | 0.3 | 0.3 |
| **C Proofreading and total reconstruction time in PWD** |  |  |  |  |
| **Minutes per false merge** | **Time component** | **All objects** | **10 objects** | **10 minutes 2D painting** |
| 1 | Proofreading | 7.1 | 8.0 | 8.8 |
|  | **Total reconstruction** | **1,007.4** | **33.3** | **9.1** |
| 3 | Proofreading | 19.2 | 21.5 | 23.2 |
|  | **Total reconstruction** | **1,019.5** | **46.8** | **23.5** |
| 10 | Proofreading | 61.7 | 68.8 | 73.6 |
|  | **Total reconstruction** | **1,062.0** | **94.1** | **73.9** |

We estimated total reconstruction time for HARRIS-15 as the sum of annotation, machine and proofreading time (Table 1). We assumed 0.1 min to fix each false split and 1, 3 or 10 min to fix each false merge. This range spans published proofreading rates^34,35^. FlyWire proofreading averaged about 1 min per edit^36^. Sparser annotation increases the proofreading burden, but the reduction in upstream annotation cuts total reconstruction time by an order of magnitude. Even when assuming the most burdensome proofreading effort of 10 min per false merge, sparse annotation provides the most efficient path to accurate data.

## Discussion

We developed a general-purpose method for generating dense 3D segmentations from sparse 2D annotations and demonstrated its applicability across imaging modalities, biological targets, and segmentation tasks. The method can process new volumetric datasets without pre-trained domain-specific models or memory-intensive computational workflows. Its 3D network never sees an image, only stacked 2D predictions, so the connectivity it learns from synthetic shapes transfers to any imaging modality or structure. Its rapidly generated segmentations were effective pseudo ground truth for iterative bootstrapping. Because its output is a dense 3D segmentation, the method is a module that supplies training data to any 3D segmentation model that learns from dense labels.

A dedicated 3D model trained using dense, representative ground truth will achieve the highest accuracy within its domain. Our method complements such specialist pipelines by generating the dense data needed to train and evaluate them. Combining training data from diverse sources improves generalization even when individual pseudo-labels contain errors^37^, and lightweight generalist models trained on combined datasets can outperform dataset-specific specialists^38^. In HARRIS-15, pseudo ground truth covering all of Volume 2 trained a better 3D model than the expert labels of a cylindrical subregion (Fig. 7), so coverage can matter more than label precision. Our method enables rapid generation of pseudo ground truth across many volumes with different imaging conditions and biological targets. Theoretical work on iterative bootstrapping demonstrates that exponentially scaling pseudo ground-truth generation maximizes model improvement per unit of compute^39^, which supports the bootstrapping strategy we use.

Foundation models still need prompting or fine-tuning for specific biological targets, so microscopy image analysis increasingly combines them with task-specific modules^5^. Our 2D→3D pipeline is complementary to segmentation foundation models; it can take their 2D outputs directly as sparse input labels. We demonstrated this with SAM-generated labels on the LICONN and EPI datasets, where 5–10 minutes of SAM-assisted annotation produced 2D labels sufficient for dense 3D segmentation. As 2D foundation models improve, our 2D→3D pipeline can use their higher-quality labels without changes. Recent work combining SAM with 3D watershed algorithms for cell structure extraction highlights the difficulty of obtaining consistent 3D reconstructions from 2D segmentations^40^. Our pipeline provides a learned solution to this problem.

The cost advantage of sparse annotation depends on volume size. Because annotation is a one-time cost and proofreading increases with object count, more accurate training data becomes preferable for sufficiently large volumes. An iterative workflow limits this proofreading cost: a subset of the bootstrapped segmentation is proofread, the corrected objects are added to the training set, and the 3D model is retrained. Each round should lower the residual error rate while keeping human effort far below that of dense annotation. Proofreading itself is becoming faster and more automated^35,41^, which further lowers the cost per edit.

Pipeline developers must decide where human effort is best spent to obtain dense annotation: painting voxels, proofreading, or supervising AI-suggested corrections. Sparse 2D annotation, automated 2D→3D inference and targeted proofreading together form one efficient allocation. The next step is to measure the marginal value of each kind of human feedback.

Predicted LSDs encode the shape the network expects at each voxel, so they disagree with the LSDs of the final segmentation wherever a merge or split contradicts that shape (Supplementary Fig. 9). Future work should use the LSD inconsistencies in targeted proofreading tools to guide users or automated systems to specific error regions^34,42–46^. Automatic error correction methods could learn to correct errors identified by LSD inconsistencies, prompting users for single-click confirmations of predicted corrections^17,45–47^. Bootstrapping can also use the LSD inconsistencies directly for targeted model refinement, with inconsistent regions weighted more heavily in the loss.

The method has three main limitations. First, the 3D network cannot infer connectivity across gaps that are ambiguous even to human annotators, as occurs with thick sections, missing sections or severe folds (Supplementary Fig. 10). Simulating missing or misaligned sections in the synthetic training stacks, analogous to the defect augmentations used for CREMI^29^, could mitigate such gaps. Second, structures absent from the sparse labels are out of distribution for the 2D network, so labels should sample the diversity of the volume. Third, bootstrapped segmentations contain errors and should be proofread before they serve as final training data for production models.

Dense voxel ground truth remains the gold standard for 3D microscopy segmentation. By reducing the total cost of generating training data, our method makes this standard affordable for laboratories with limited resources. Our bootstrapper command-line interface is fully scriptable, so an AI agent could run the whole bootstrapping loop: selecting annotations, training, inference, evaluation and iteration.

## Methods

### Sparsity generation and manual annotation

To evaluate the impact of annotation density on model performance, we generated varying levels of sparse training data for Volume 1 of each dataset. We performed instance-level ablations to create subsets containing 1, 2, 2 adjacent, 5, 10, 50, or 100 randomly selected objects. Each level had three mutually exclusive random selections. We also explored disk ablations on top of instance ablations to simulate imprecise brush-stroke annotations (Supplementary Fig. 1; Supplementary Note 1). To assess usability in practice, we collected 30 minutes of non-expert sparse 2D annotations on a single section for each dataset (Supplementary Fig. 2) and partitioned them into three subsets representing 10-minute annotation budgets.

### Network architectures

We used 2D and 3D U-Net architectures to learn dense cell boundaries, local shape descriptors (LSDs), or both. Voxel-wise direct-neighbor affinities were used to represent boundaries, capturing the connectivity between every voxel and its immediate neighbors^48^. This representation resolves ambiguities when partitioning voxels into instances that share boundaries or lack explicit background separation. We use the 3D multitask LSD (MTLSD) model, trained on dense ground truth, as our 3D baseline^29^ (Figs. 2 and 7; Statistics and reproducibility). Detailed network architectures, training pipelines, and parameter values are provided in Supplementary Notes 2, 3 and Supplementary Tables 3, 4. The lightweight design of the algorithms enables generation of 3D segmentations on standard consumer hardware without cluster access. High-performance computing resources (TACC’s Lonestar6) were used for large-scale grid searches.

### 2D→3D

The first stage transforms sparse 2D annotations into dense 2D predictions. A 2D U-Net is trained using the sparse annotations to predict dense 2D affinities, LSDs, or both. During training, random fields of view are sampled from annotated sections with standard augmentations (Supplementary Note 3; Supplementary Tables 3, 4). A masked loss function restricts supervision to labeled regions, preventing the network from learning background features from unannotated instances^21^. During inference, the trained 2D network is applied section-wise to the target volume, producing a stack of dense 2D predictions.

The second stage transforms the stack of independent 2D predictions into a coherent 3D segmentation by inferring z-axis connectivity. A lightweight 3D U-Net pre-trained entirely using synthetic 3D labeled data (Fig. 3c) maps stacked 2D predictions to 3D boundaries. Because the network learns geometric priors of 3D connectivity from synthetic shapes, it does not require manual 3D annotation from the target dataset. During training, the stacked 2D predictions computed from the synthetic labels are heavily augmented to allow the network to generalize to true stacked 2D predictions.

### Synthetic 3D label generation

Synthetic 3D training data was generated using three distinct strategies to simulate diverse biological structures. For each batch, we drew an anisotropy factor *a* from 1 to the integer part of the axial-to-lateral voxel size ratio (1 to 6 for HARRIS-15). We generated labels *a* times taller along z and then subsampled them by *a*. In the first strategy, a random number of voxels *N* ∈ [25,50] (scaled to [25, 50*a*] for the anisotropy factor *a*) within a zero-initialized array are set to 1, uniquely labeled, and grown outward *D* ∈ [25,40] voxels without overlap. The background is assigned a unique non-zero ID. The second strategy extends this by dilating a speckled binary array section-wise using either a 3×3 square or a disk structuring element with a random radius (*r* ∈ [1,5]).

Foreground instances are uniquely labeled, and objects are expanded into unoccupied space using a distance transform of the background (Fig. 3d). The third strategy generates organic shapes by applying a Gaussian filter (*σ* = 5) to an array of random float values to identify peaks. A maximum filter accentuates these peaks. A watershed algorithm uses them as seeds and grows them (Fig. 3d). A fourth ensemble strategy, which was an equal mix of the above three strategies, was also included. The implementation is available as a Gunpowder node that generates the labels during training.

### Post-processing

For the ablation and bootstrapping experiments (Figs. 2, 6 and 7), dense 3D instance segmentations were generated from predicted affinities using a seeded watershed-based agglomeration pipeline^49–51^. First, predicted affinities were thresholded to create a binary mask, from which a distance transform was computed to identify local maxima. These maxima seeded a watershed algorithm, resulting in an initial over-segmentation of supervoxels. A region adjacency graph (RAG) was constructed where each node represented a supervoxel center of mass, and edges connected touching supervoxels. During final agglomeration, we hierarchically merged edges in the RAG in order of decreasing affinity weight until a stopping criterion was met. For the EPI comparison, the additional datasets and the bootstrapping-path comparison (Figs. 4 and 5; Supplementary Fig. 6), mutex watershed instead produced the segmentations from the predicted affinities^52^.

### Evaluation metrics

We assessed segmentation accuracy using normalized VOI and the min-cut metric (MCM). VOI measures the information-theoretic dissimilarity between segmentation and ground truth, quantifying split and merge errors^26,27^. MCM approximates human proofreading effort as the number of split and merge edit operations required to match ground-truth skeletons^29^. We automatically generated ground-truth evaluation skeletons from ground-truth labels by computing a minimum spanning tree of supervoxel centers of mass. We obtained the supervoxels by watershed over-segmentation of ground-truth affinities. Ground-truth labels were filtered to remove objects smaller than 500 pixels.

### Comparison with Cellpose and uSegment3D

The comparison (Fig. 4) used the volume of the uSegment3D tutorial^13^, taken from the validation split of the ovules dataset^25^ (Supplementary Table 1). This volume is not one of the two EPI volumes of the ablation. Cellpose^10^ (model cyto; uSegment3D 0.1.4) ran inside uSegment3D on all three orthogonal views of the volume. uSegment3D fused the resulting probability and flow maps into a 3D consensus segmentation with the published ovule tutorial script. We changed only the input and output paths^53^. Before scoring, label boundaries of the uSegment3D output were eroded as in the ground truth. For Cellpose + uSegment3D, Fig. 4b lists the labeled voxels, objects and images of the training set of the Cellpose cyto model.

For the 2D→3D method, we trained three 2D U-Nets on labels from this same volume and evaluated them on it. Their training labels were the dense ground truth, a single ground-truth section, or sparse SAM^7^ labels drawn on three sections in about 5 minutes. We drew these labels with an interactive Neuroglancer tool running the Segment Anything automatic mask generator (ViT-L). Each 2D network predicted 2D LSDs, and the same 3D network of the 2D→3D method, trained only on synthetic labels, converted them to 3D affinities. Mutex watershed produced the segmentation. For each variant, the watershed bias was selected from a grid of 25 settings by the score against the ground truth. The annotation-free baseline applied an Otsu threshold to the raw image, then a watershed on the distance transform of the cell interiors. Accuracy was normalized VOI (sum)^26,27^ against the ground truth over the full volume.

### Additional datasets

The five additional datasets (Fig. 5) were processed with the same pipeline. Imaging parameters, sources, and training labels are listed in Supplementary Table 1. The labels that trained the 2D network were sections of the published ground truth, except for LICONN, where they were drawn with an interactive SAM tool. Mutex watershed produced the segmentation from the predicted 3D affinities.

LICONN^30^: neuron instance segmentation in an expansion-microscopy volume of mouse hippocampal CA1 (ExPID82). Six sections in six crops were labeled in about 10 minutes with the interactive SAM tool of the bootstrapper repository (Code availability). The 2D U-Net predicted 2D LSDs and 2D affinities; the LSDs were the input of the 3D network. The evaluation crop contained one of the six training sections.

CREMI-C^24^ synaptic clefts: synaptic cleft instance segmentation in the serial-section TEM volume of *Drosophila* neuropil (sample C). We trained the 2D U-Net on two sections of the published cleft ground truth to predict 2D LSDs and 2D affinities. The 3D network used the LSDs as input. The evaluation crop contained one of the two training sections.

MitoEM-H^31^: mitochondria instance segmentation in the multi-beam SEM volume of layer II of adult human frontal cortex. We trained the 2D U-Net on two sections of the published ground truth to predict 2D LSDs and 2D affinities. The 3D network used both as input. The evaluation crop lay outside the training sections.

PRISM^32^: neuron instance segmentation in an 18-channel expansion light microscopy volume of mouse hippocampus (E11 Bio open data, sample 6). We trained the 2D U-Net on ten contiguous sections of the published proofread labels, using all 18 channels. The network predicted 2D LSDs and 2D affinities as inputs to the 3D network. The 2D→3D method segmented the full volume; the section shown in Fig. 5d is one of the ten training sections.

Fluo-C2DL-Huh7^33^: cell instance segmentation over time (2D+t) in training sequence 01 of the Cell Tracking Challenge dataset (live-cell fluorescence time-lapse imaging), with time as the third axis. We trained the 2D U-Net on two frames of the published SEG ground truth to predict 2D LSDs and 2D affinities. The 3D network used the affinities as input. The comparison used the eight annotated frames, which include the two training frames.

### Bootstrapping experiment

The bootstrapping experiment (Fig. 6; Supplementary Note 4) was run for each dataset, sparsity level and repetition as a grid search over network variants, training iterations and post-processing parameters. In Stage 1, a 2D U-Net trained on the sparse 2D labels of Volume 1 predicted 2D affinities, 2D LSDs or both (2D MTLSD) at checkpoints between 5,000 and 20,000 iterations. This gave four prediction types: 2D affinities, 2D LSDs, and each of the two from the 2D MTLSD network. A 3D network of the 2D→3D method, trained on synthetic labels from one of the four generation strategies, inferred 3D affinities at checkpoints between 5,000 and 20,000 iterations. We then applied seeded watershed with hierarchical agglomeration over a grid of parameters (Supplementary Table 5). All candidate segmentations of Volume 2 were evaluated by normalized VOI and MCM, and the best one became the pseudo ground truth for Stage 2. In Stage 2, we trained a 3D MTLSD network (Supplementary Table 6) on this unproofread pseudo ground truth. Using checkpoints at 20,000, 35,000 and 50,000 iterations, we predicted 3D affinities and LSDs of Volume 1. We applied post-processing over the grids in Supplementary Tables 7, 8. Statistics and reproducibility describes how thresholds and checkpoints were selected. The non-bootstrapped baseline was a 3D MTLSD network trained on the dense ground truth of Volume 2 of the same repetition and evaluated on Volume 1.

### Proofreading time estimate

Total reconstruction time for HARRIS-15 (Table 1) was the sum of manual annotation time, machine time and estimated proofreading time, in person-workdays (PWD; 1 PWD = 5 h). Proofreading time was 0.1 min per false split and 1, 3 or 10 min per false merge, applied to the edit counts of Table 1. K.M.H. estimated the dense annotation time (1,000 PWD) from the original expert annotation of HARRIS-15^22^, at 5 h of annotation per workday. Annotation of 10 objects was set to 25 PWD, from an estimate by K.M.H. of 2.5 PWD per object. Ten minutes of painting was set to 0.03 PWD. Machine time was 0.3 PWD for all conditions.

### Statistics and reproducibility

For the simulated sparsity levels, we drew objects in three mutually exclusive random selections per level. We split each 30-minute painting into three 10-minute subsets (Supplementary Note 1). For the all-objects and 30-minute conditions, the three repetitions differ only in the random initialization of the network weights and the random seeds of training. Fig. 2, Fig. 7 and Table 1 report descriptive statistics only. In Fig. 2 and Supplementary Figs. 3–5, Δ is the 2D→3D segmentation score minus the 3D MTLSD baseline score. The baseline was trained on the dense ground truth of the same repetition. Points are means and bars ±1 s.d. over n = 12 differences (4 prediction types × 3 repetitions). The four prediction types of a repetition share one baseline model, so the 12 differences are not independent. For each prediction type and repetition, we selected the grid configuration with the best value of the plotted metric. The VOI sum, split and merge panels can therefore use different configurations. In Fig. 7, Supplementary Figs. 7, 8 and Table 1, we selected the merge threshold with the fewest total edits for each post-processing configuration. For the VOI merge panel of Supplementary Fig. 8, we instead selected the threshold with the lowest VOI sum. Then the checkpoint with the best value of each reported metric was selected per bootstrapped model. Table 1 counts are means over the same 12 models, so its rows can come from different checkpoints. Points are means and bars ±1 s.d. over n = 12 bootstrapped models (4 pseudo ground-truth network variants × 3 repetitions). We computed Δ relative to the non-bootstrapped 3D MTLSD trained on the dense ground truth of the same repetition. Supplementary Fig. 6 shows six points, one per bootstrapping-path pair, without error bars. Supplementary Note 5 reports the paired differences and a two-sided sign test.

## Data availability

The image volumes, their ground-truth labels and a sparse 2D annotation for each dataset are available on Zenodo at https://doi.org/10.5281/zenodo.21223591. The download_data.sh script of github.com/ucsdmanorlab/bootstrapper_nm_capsule downloads these data. The original sources of all datasets are listed in Supplementary Table 1.

## Code availability

The bootstrapper framework is open source at github.com/ucsdmanorlab/bootstrapper. Its command-line interface has modules for data preparation, 2D→3D and 3D models, training, blockwise inference and post-processing, evaluation and error identification. The current release is version 1.0.0 (github.com/ucsdmanorlab/bootstrapper/releases/tag/v1.0.0). The napari plugin for the 2D→3D method is at github.com/ucsdmanorlab/napari-bootstrapper. The interactive SAM annotation tool is ng_interactive_sam.py in the bootstrapper repository. The code that reruns the experiments with version 1.0.0, with a Dockerfile and pinned requirements, is at github.com/ucsdmanorlab/bootstrapper_nm_capsule. The results in this paper were produced with Bootstrapper v0.3.2 (github.com/ucsdmanorlab/bootstrapper/releases/tag/v0.3.2). A companion page with Neuroglancer views of the segmentations is at ucsdmanorlab.github.io/sparsity.

## Acknowledgements

U.M. is supported by NIA P30AG068635 (Nathan Shock Center), the David F. and Margaret T. Grohne Family Foundation, Core Grant application NCI CCSG (CA014195), NIDCD R01DC021075-01, NSF NeuroNex Award (2014862), the L.I.F.E. Foundation, and the CZI Imaging Scientist Award DOI https://doi.org/10.37921/694870itnyzk from the Chan Zuckerberg Initiative DAF, an advised fund of Silicon Valley Community Foundation (funder DOI 10.13039/100014989). K.M.H. and V.V.T. are supported by NIH R01MH095980, NSF NeuroNex Technology Hub Award (1707356), NSF NeuroNex Award (2014862), and NSF NCS Award (2219864).

The authors thank all the manual annotators for their valuable work. Special acknowledgment to Patrick H. Parker for expert curation and proofreading of the HARRIS-15 dataset. The authors also acknowledge James Carson and the Texas Advanced Computing Center (TACC) at The University of Texas at Austin for providing HPC resources that have contributed to the research results reported within this paper. URL: http://www.tacc.utexas.edu

## Author contributions

V.V.T. and A.S. conceived the method and developed its first prototype. V.V.T. wrote the bootstrapper software, the napari plugin and the annotation tool, designed and ran the experiments, and wrote the manuscript. K.M.H. and U.M. supervised the project. All authors edited the manuscript.

