## Supplementary Information for "A general method for bootstrapping dense 3D segmentations from sparse 2D annotations"

#### Supplementary Note 1. Annotation sparsity definitions

We define a *sparsity* as an amount of manual annotation that we use to train networks with a weighted loss function to infer dense predictions. We varied annotation density at the object and sub-object levels to characterize the minimum annotation required for the 2D→3D method to produce usable segmentations.

*Object-level sparsities.* We subsampled the number of annotated objects in the training volume at eight levels: 1, 2, 2 adjacent, 5, 10, 50, 100, and all objects (dense) (Supplementary Fig. 1, top row). For each level, three mutually exclusive random selections of the objects were made to provide independent *repetitions*. For all objects and the full 30-minute annotations below, the three repetitions differ only in random weight initialization and random training seeds.

*Disk sparsities*. To further reduce annotation within selected objects, we centered fixed-radius circular masks at randomly sampled points within the field of view (FOV). We retained only the label regions intersected by these disks (Supplementary Fig. 1, bottom row). This simulates rapid brush-stroke annotations, in which an annotator paints partial labels across object boundaries and interiors without fully tracing each object. Three disk sparsity levels (1, 2, 3 disks) were combined with each object-level sparsity, yielding 32 total sparsity conditions.

*Non-expert manual annotations.* In addition to simulated sparsities, we generated non-expert 2D annotations in approximately 30 minutes per dataset for CREMI[1], EPI[2], FIB-25[3], and HARRIS-15[4] datasets (Supplementary Fig. 2). These annotations were split into three approximately 10-minute subsets to assess the effect of annotation time on segmentation quality.

We based disk sparsities on the structure of local shape descriptors (LSDs)[5]: the most informative regions for learning LSDs are object boundaries and interiors. Partial annotations that sample both regions may therefore retain sufficient signal for training. We observed no systematic prediction artifacts from disk-masked labels, despite incorrect LSD targets near disk centers where smooth gradients replace sharp boundaries. The absence of artifacts is likely due to the regression behavior of the network in ambiguous regions. There, predictions converge toward intermediate values that approximate the correct LSD values for non-boundary regions.

#### Supplementary Note 2. Network architectures

All networks were trained using Gunpowder (github.com/funkelab/gunpowder) and PyTorch with U-Net architectures from funlib.learn.torch (github.com/funkelab/funlib.learn.torch). Code for

LSDs is available at github.com/funkelab/lsd. Code for the 2D→3D and bootstrapping pipelines is available at github.com/ucsdmanorlab/bootstrapper and github.com/ucsdmanorlab/bootstrapper_nm_capsule.

**2D U-Nets (2D→3D method).** The 2D U-Nets[6] consisted of three layers with downsampling factors of 2, 2 per layer. Twelve initial feature maps were used with a multiplication factor of 6 between layers. A final convolution and sigmoid activation produced 2 (affinities), 6 (LSDs), or 8 (multitask LSD, MTLSD) output channels. Network and training parameters are listed in Supplementary Table 3.

**3D U-Net (2D→3D method).** The lightweight 3D U-Net[7] in the 2D→3D pipeline consisted of two layers with downsampling factors of 1, 2, 2. The first downsampling and last upsampling layers used kernels of size 2, 3, 3, while all other layers used 1, 3, 3 kernels. Five initial feature maps were used with a multiplication factor of 5 between layers. The network accepts stacked 2D predictions rather than raw images and produces 3 output channels (3D affinities) after sigmoid activation. We trained the 3D networks of the 2D→3D method only on synthetic labels, without using target data (Supplementary Table 4).

**3D MTLSD network (3D baseline).** The 3D MTLSD[5] network used for bootstrapping was a 3D U-Net with two separate decoder heads for affinities and LSDs. It had three layers with downsampling factors of 1, 2, 2, and used 2D convolution kernels in the bottleneck and adjacent layers. Thirteen initial feature maps were used with a multiplication factor of 6 between layers. Each decoder head produced 3 (affinities) or 10 (LSDs) output channels after sigmoid activation (Supplementary Table 6).

**Computational requirements**. The number of trainable parameters and training GPU memory usage for each network are listed in Supplementary Table 2. All 2D→3D networks require less than 3 GB of GPU memory, which enables training on consumer-grade hardware.

### Supplementary Note 3. Training pipelines

For all sparse 2D and 3D networks and the bootstrapping 3D MTLSD network, we randomly sampled each batch from the available sections or volume. For each batch, the raw data was first normalized and padded with zeros. Labels were padded with the maximum padding required to contain at least 0.01% of labeled ground-truth data within the image, assuming a worst-case rotation of 45 degrees. For 3D MTLSD, the threshold was 10%, since more pseudo ground-truth labels are available. Data was randomly sampled and augmented with elastic transformations, random mirrors and transposes, Gaussian blur, and intensities (see Supplementary Tables 3 and 6 for augmentation hyper-parameters used for all networks). For the networks with affinities as an output, a scale array was created to balance loss between target affinity labels.

For the lightweight 3D networks in the 2D→3D method, each training batch begins as a 3D array of zeros. Synthetic 3D labels are randomly grown using the strategies illustrated in Fig. 3d. We augmented the labels with elastic transformations and random mirrors and transposes, then used them to simulate stacked 2D affinities or LSDs. The stacked 2D affinities or LSDs were then augmented with random noise, intensities and Gaussian blur to simulate realistic stacked

2D predictions. Finally, target 3D affinities were computed, and a scale array was created to balance loss between class labels.

### Supplementary Note 4. Bootstrapping experiment procedure

The bootstrapping experiment (Fig. 6) used a grid search over network variants, training iterations, and post-processing parameters. We ran this grid search for each combination of dataset, sparsity level, and repetition. Dataset details, including imaging modality, resolution, volume sizes, and number of annotated objects, are provided in Supplementary Table 1.

**Stage 1: 2D→3D segmentation of Volume 2**.

1. A 2D U-Net was trained using sparse 2D annotations of Volume 1. Three network variants were explored, differing by output representation: 2D affinities, 2D LSDs, or both (2D MTLSD).
2. Trained 2D U-Net checkpoints at iterations {5,000; 10,000; 15,000; 20,000} generated stacked 2D predictions. The four prediction types were: 2D affinities, 2D LSDs, 2D affinities (from MTLSD), and 2D LSDs (from MTLSD).
3. A 3D U-Net was trained on synthetic 3D labels from one of four label generation strategies (Fig. 3d). It inferred 3D affinities from the stacked 2D predictions at iterations {5,000; 10,000; 15,000; 20,000}.
4. Post-processing was applied in a grid search over watershed and agglomeration parameters (Supplementary Table 5) to generate candidate 3D segmentations of Volume 2.
5. All candidate segmentations were evaluated using normalized variation of information (VOI) and the min-cut metric (MCM)[5]. The best-performing segmentation became the pseudo ground-truth for Stage 2.

**Stage 2: 3D segmentation of Volume 1.**

1. A 3D MTLSD network was trained using the unproofread pseudo ground-truth segmentation of Volume 2 from Stage 1.
2. Trained 3D MTLSD checkpoints at iterations {20,000; 35,000; 50,000} predicted 3D affinities and LSDs of Volume 1.
3. Post-processing was applied in a grid search (Supplementary Tables 7, 8) to generate candidate 3D segmentations of Volume 1.
4. All candidate segmentations of Volume 1 were evaluated against dense ground-truth annotations. The best-performing segmentation became the representative bootstrapped result for that dataset, sparsity, pseudo ground-truth network variant and repetition.

The total parameter grid sizes are reported in Supplementary Tables 5, 7, and 8. Quantitative results across all sparsities are shown for the 2D→3D method in Supplementary Figs. 3–5 and for bootstrapped 3D segmentations in Supplementary Figs. 7, 8.

### Supplementary Note 5. Comparing bootstrapping paths

In the bootstrapping experiment (Fig. 6), a chain of two models starts and ends with Volume 1. A first model, trained on labels of Volume 1, segments Volume 2. A second model, trained on that unproofread segmentation, segments Volume 1. The final segmentation could benefit from volume-specific biases propagated through the chain. Such biases are acceptable for bootstrapping and iterative refinement, but we measured their effect by comparing two bootstrapping paths.

**Experiment.** We cut the HARRIS-15 Apical volume into three z-slabs, A (sections 1–34), B (35–68) and C (69–101), each 901 × 928 pixels (Supplementary Table 1). For a target evaluation slab V1, we compared (1) a direct path (V1→V2→V1) and (2) a control path (V3→V2→V1). In the direct path, the first model is trained on the dense labels of V1 itself; in the control path, it is trained on the dense labels of the independent third slab V3. In both paths the first model is a 3D MTLSD network trained for 30,000 iterations on the dense labels of its source slab; it segments the intermediate slab V2. The second model is a 3D MTLSD network trained for 30,000 iterations on that unproofread segmentation; it segments V1. We scored each final segmentation against the dense labels of V1 by normalized VOI (sum), taking the minimum over a grid of 25 post-processing thresholds. The labels of V1 contain V1-specific statistics. If these statistics propagated through the chain as meaningful bias, the direct path should systematically outperform the control path.

**Results.** We ran this comparison for all six ordered pairs of the three slabs (Supplementary Fig. 6). The direct path scored better in 3 of 6 pairs. The difference in normalized VOI (sum), direct minus control, had a mean of −0.0024 (s.d. 0.0159) over the six pairs (two-sided sign test, $p = 1.0$). We found no systematic advantage of the direct path, although the six pairs reuse the same three slabs and are not independent.

| Dataset name and link | Imaging modality | Voxel size (nm, ZYX) | Content | Volume or crop name | Size (pixels, ZYX) | Objects | Training labels |
|---|---|---|---|---|---|---|---|
| HARRIS-15[4] / neurodata.io/data/kharris15 | TEM | (50, 2, 2), downscaled to (50, 8, 8) | *Rattus* hippocampal neuropil | Apical | (101, 900, 927) | 444 | Dense GT (and generated sparsities/ablations) |
| | | | | Oblique | (70, 674, 504) | 181 | |
| FIB-25[3] / janelia.org/tools-and-data-release | FIBSEM | (8, 8, 8) | *Drosophila* optic medulla neuropil | tstvol-520-1 | (520, 520, 520) | 1690 | |
| | | | | tstvol-520-2 | (520, 520, 520) | 2031 | |
| CREMI[1] / cremi.org/data | SSTEM | (40, 4, 4), downscaled to (40, 8, 8) | *Drosophila* calyx neuropil | A [:62] | (62, 625, 625) | 546 | |
| | | | | A [62:] | (63, 625, 625) | 647 | |
| | | | *Drosophila* axon tract | B [62:] | (62, 625, 625) | 450 | |
| | | | | B [:62] | (63, 625, 625) | 581 | |
| | | | *Drosophila* axon tract | C [:63] | (63, 625, 625) | 333 | |
| | | | | C [63:] | (62, 625, 625) | 312 | |
| EPI[2] / osf.io/8jz7e | LM | (235, 75, 75) | *Arabidopsis* ovule cells | N_294 | (318, 960, 953) | 3333 | |
| | | | | N_511 | (248, 791, 667) | 1053 | |
| EPI[2], uSegment3D tutorial image[8] | LM | (235, 235, 235), after isotropic resampling | | N_464 | (540, 733, 391) | 1686 | Dense GT; one GT section (z = 270); or SAM labels on three sections (z = 107, 270, 392) |
| LICONN[9] / gs://liconn-public/ExPID82_1 | Expansion LM | (24, 18, 18) | Mouse hippocampal CA1 neuropil | crop_5 | (25, 512, 513) | 371 | Six sections in six crops, SAM-assisted (about 10 min) |
| CREMI-C clefts[1] / cremi.org/data | SSTEM | (40, 4, 4) | *Drosophila* synaptic clefts (sample C) | crop_sample_c | (75, 1024, 1024) | 85 | Two sections of the published GT (z = 108, 141) |
| MitoEM-H[10] / mitoem.grand-challenge.org | Multi-beam SEM | (30, 8, 8) | Human frontal lobe (layer II) mitochondria | crop_mito | (75, 1024, 1024) | 149 | Two sections of the published GT (z = 19, 95), outside the evaluation crop (z = 178–252) |
| PRISM[11] / s3://e11bio-prism | Expansion LM, 18 channels | (400, 168, 168) | Mouse hippocampal neurons (protein barcodes) | instance_example (sample 6) | (200, 500, 500) | 221 | Ten sections of the published labels (z = 540–549) |
| Fluo-C2DL-Huh7[12] / celltrackingchallenge.net/2d-datasets | Live-cell fluorescence time-lapse (2D+t) | 650 × 650 nm per pixel (y, x); 15 min per frame | Huh7 cells expressing YFP-TIA-1 | Sequence 01 | (30 frames, 1024, 1024) | 253 | Two frames of the published SEG GT (frames 2, 6) |

**Supplementary Table 1 | Overview of datasets.** We applied a size filter of 500 pixels when counting objects, except in the Fig. 4 comparison volume, where we counted all labeled objects. Voxel sizes are the imaged sizes, except for the Fig. 4 comparison volume, which uses the nominal size after resampling, and LICONN, whose size is expansion corrected. The rows for the five generalization datasets (Fig. 5) and the Fig. 4 comparison volume list the evaluation volumes. Their last column lists training labels for the 2D U-Net, with z values indicating section indices in the source dataset. For the six ablation datasets, the training labels are the annotation levels of Supplementary Note 1. For Fluo-C2DL-Huh7 the count is cell-frame instances over the eight annotated frames. The bootstrapping-path comparison (Supplementary Note 5) uses three z-slabs of the Apical volume: A (sections 1–34), B (35–68) and C (69–101), each 901 × 928 pixels. TEM, transmission electron microscopy; FIBSEM, focused ion beam scanning electron microscopy; GT, ground truth; SSTEM, serial-section TEM; LM, light microscopy; SEM, scanning electron microscopy; YFP, yellow fluorescent protein; ZYX, axis order z, y, x.

| Network | Number of trainable parameters | Training GPU memory |
|---|---|---|
| 2D U-Net (Affinities) | 88,886,116 | 2.8 GB |
| 2D U-Net (LSDs) | 88,886,220 | 2.5 GB |
| 2D→3D U-Net | 264,206 | 0.6 GB |
| 3D U-Net (MTLSD) | 187,077,748 | 6.0 GB |

**Supplementary Table 2 | Number of trainable parameters and training GPU memory for the tested models.**

| Parameter | Value |
|---|---|
| Input feature maps | 12 |
| Layer feature map scale | 6 |
| Downsampling factors | [[2, 2], [2, 2], [2, 2]] |
| Input shape | [196, 196] |
| Output shape | [104, 104] |
| Loss | Weighted MSE |
| Optimizer | Adam |
| Learning rate | $0.5 \times 10^{-4}$ |
| β1 | 0.9 |
| β2 | 0.999 |
| ε | $1 \times 10^{-8}$ |
| Iterations | 20,000 |

| Augmentation | Parameter | Value |
|---|---|---|
| Elastic | Control point spacing | (8, 8) |
| | Jitter sigma | (2, 2) |
| | Subsample | 4 |
| Rotation | Axis | x, y |
| | Angle | in $[0, 2\pi]$ |
| Mirror | Axes | x, y |
| Transpose | Axes | x, y |
| Noise | Mode | Gaussian |
| Intensity | Scale | in [0.9, 1.1] |
| | Shift | in [−0.1, 0.1] |
| Blur | Sigma | in [0.0, 1.5] |

**Supplementary Table 3 | 2D networks and training parameters for HARRIS-15.** Top: Network architecture and training parameters. Bottom: Data augmentation parameters.

| Parameter | Value |
| --- | --- |
| Input feature maps | 5 |
| Layer feature map scale | 5 |
| Downsampling factors | [[1, 2, 2], [1, 2, 2]] |
| Kernel sizes down | [[2, 3, 3], [2, 3, 3]],<br>[[1, 3, 3], [1, 3, 3]],<br>[[1, 3, 3], [1, 3, 3]] |
| Kernel sizes up | [[1, 3, 3], [1, 3, 3]],<br>[[2, 3, 3], [2, 3, 3]], |
| Input shape | [10, 148, 148] |
| Output shape | [6, 108, 108] |
| Loss | Weighted MSE |
| Optimizer | Adam |
| Learning rate | $0.5 \times 10^{-4}$ |
| β1 | 0.9 |
| β2 | 0.999 |
| ε | $1 \times 10^{-8}$ |
| Iterations | 20,000 |

**Supplementary Table 4 | Network architecture and training parameters of the 3D networks in the 2D→3D pipeline.**

| Parameter | Values | Number of variations |
|---|---|---|
| Synthetic 3D labels generation | Strategies A, B, C, (A+B+C) | 4 |
| 3D affinities iteration | {5000, 10000, 15000, 20000} (2D U-Net)<br>x<br>{5000, 10000, 15000, 20000} (3D U-Net) | 16 |
| Rep | rep_1, rep_2, rep_3 | 3 |
| Sparsity | 10min_paint_{2d,3d}<br>+<br>paint_{2d, 3d}<br>+<br>{disk_{0,1,2,3}} x<br>obj_{001,002,002a,005,010,100,dense}<br>+<br>3D_dense | 33 |
| Watershed minimum seed distance | 10 | 1 |
| Hierarchical merge function | “mean”, “hist_quant_50”, “hist_quant_75” | 3 |
| Watershed merge thresholds | 0.3, 0.35, 0.4, …, 0.75, 0.8 | 11 |
| Total parameter grid size | 209,088 | |

**Supplementary Table 5 | Parameter grid explored for generating 2D→3D segmentations for HARRIS-15.**

| Parameter | Value |
|---|---|
| Input feature maps | 13 |
| Layer fmap scale | 6 |
| Downsampling factors | [[1, 2, 2], [1, 2, 2], [1, 2, 2]] |
| Kernel sizes down | [[3, 3, 3], [3, 3, 3]], [[3, 3, 3], [3, 3, 3]],<br>[[1, 3, 3], [1, 3, 3]], [[1, 3, 3], [1, 3, 3]] |
| Kernel sizes up | [[1, 3, 3], [1, 3, 3]],<br>[[3, 3, 3], [3, 3, 3]], [[3, 3, 3], [3, 3, 3]] |
| Input shape | [20, 196, 196] |
| Output shape | [4, 104, 104] |
| Loss | Weighted MSE |
| Optimizer | Adam |
| Learning rate | $0.5 \times 10^{-4}$ |
| β1 | 0.9 |
| β2 | 0.999 |
| ε | $1 \times 10^{-8}$ |
| Iterations | 50,000 |

| Augmentation | Parameter | Value |
|---|---|---|
| Elastic | Control point spacing | (8, 50, 50) |
| | Jitter sigma | (0, 2, 2) |
| | Subsample | 4 |
| | Scale interval | (0.75, 1.25) |
| Rotation | Axis | x, y |
| | Angle | in $[0, 2\pi]$ |
| Mirror | Axes | x, y, z |
| Transpose | Axes | x, y |
| Noise | Mode | Gaussian |
| Intensity | Scale | in [0.9, 1.1] |
| | Shift | in [−0.1, 0.1] |
| Blur | Sigma | in [0.0, 1.5] |

**Supplementary Table 6 | Baseline 3D networks and training parameters for HARRIS-15.** Top: Network architecture and training parameters. Bottom: Data augmentation parameters.

| Parameter | Values | Number of variations |
|---|---|---|
| Predicted affinities iteration | 20000, 35000, 50000 | 3 |
| Rep | rep_1, rep_2, rep_3 | 3 |
| Pseudo ground-truth networks | Affinities,<br>LSDs,<br>Affinities from MTLSD,<br>LSDs from MTLSD | 4 |
| Pseudo ground-truth sparsities | 10min_paint_{2d,3d}<br>+<br>paint_{2d, 3d}<br>+<br>obj_{001,002,002a,005,010,100} + dense | 11 |
| Normalize affinities | False | 1 |
| Watershed minimum seed distance | 10 | 1 |
| Watershed boundary mask | True | 1 |
| Hierarchical merge function | “mean” | 1 |
| Watershed merge thresholds | 0.3, 0.35, 0.4, …, 0.75, 0.8 | 11 |
| Total parameter grid size | | 4356 |

**Supplementary Table 7 | Parameter grids explored for computing VOI and MCM with bootstrapped segmentations from the 3D baseline model for HARRIS-15.**

| Parameter | Values | Number of variations |
|---|---|---|
| Predicted affinities iteration | 20000, 35000, 50000 | 3 |
| Rep | rep_1, rep_2, rep_3 | 3 |
| Pseudo ground-truth networks | Affinities,<br>LSDs,<br>Affinities from MTLSD,<br>LSDs from MTLSD | 4 |
| Pseudo ground-truth sparsities | 10min_paint_{2d,3d}<br>+<br>paint_{2d, 3d}<br>+<br>obj_{001,002,002a,005,010,100} + dense | 11 |
| Normalize affinities | False, True | 2 |
| Watershed minimum seed distance | 5, 10, 15, 20 | 4 |
| Watershed boundary mask | True, False | 2 |
| Hierarchical merge function | 'hist_quant_10', 'hist_quant_25', 'hist_quant_50',<br>'hist_quant_75', 'hist_quant_90', 'mean' | 6 |
| Watershed merge thresholds | 0.3, 0.35, 0.4, …, 0.75, 0.8 | 11 |
| Total parameter grid size | | 418,176 |

**Supplementary Table 8 | Parameter grids explored for computing VOI only with bootstrapped segmentations from the 3D baseline model for HARRIS-15.**

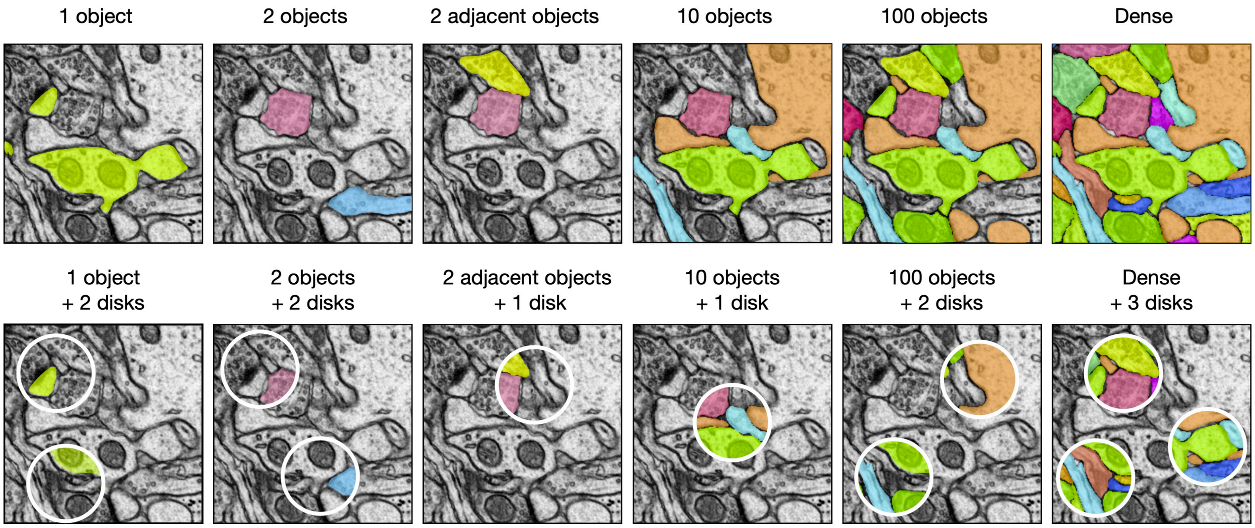


**Supplementary Fig. 1 | Example simulated sparsity regimes.** Example HARRIS-15 image with ground-truth labels as they appear in training batches (Supplementary Note 1). Top row, object-level sparsities: 1 object, 2 objects, 2 adjacent objects, 10 objects, 100 objects and all objects (dense). Bottom row, the same selections after disk masking with 1, 2 or 3 disks (white circles); only the labels inside the disks are kept. Illustration only; no quantification.

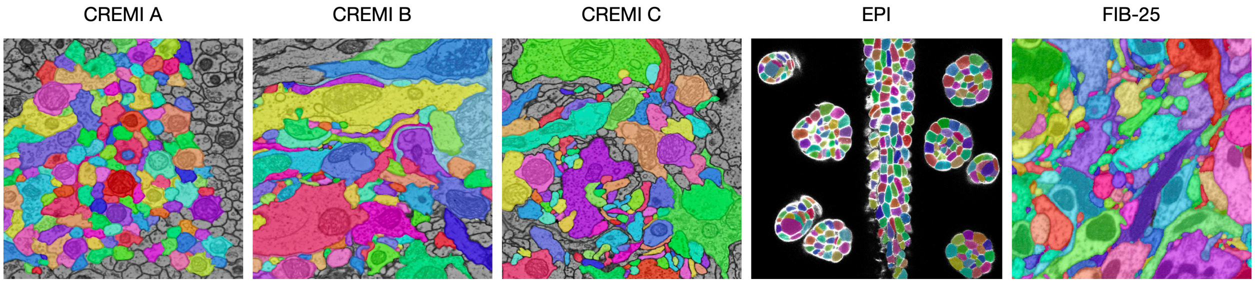


**Supplementary Fig. 2 | Thirty minutes of non-expert 2D annotations.** Example images with the overlaid non-expert annotations made in about 30 minutes for CREMI-A, -B, -C, EPI and FIB-25 (HARRIS-15 in Fig. 1b). Each 30-minute set was split into three approximately 10-minute subsets (Supplementary Note 1). Illustration only; no quantification.

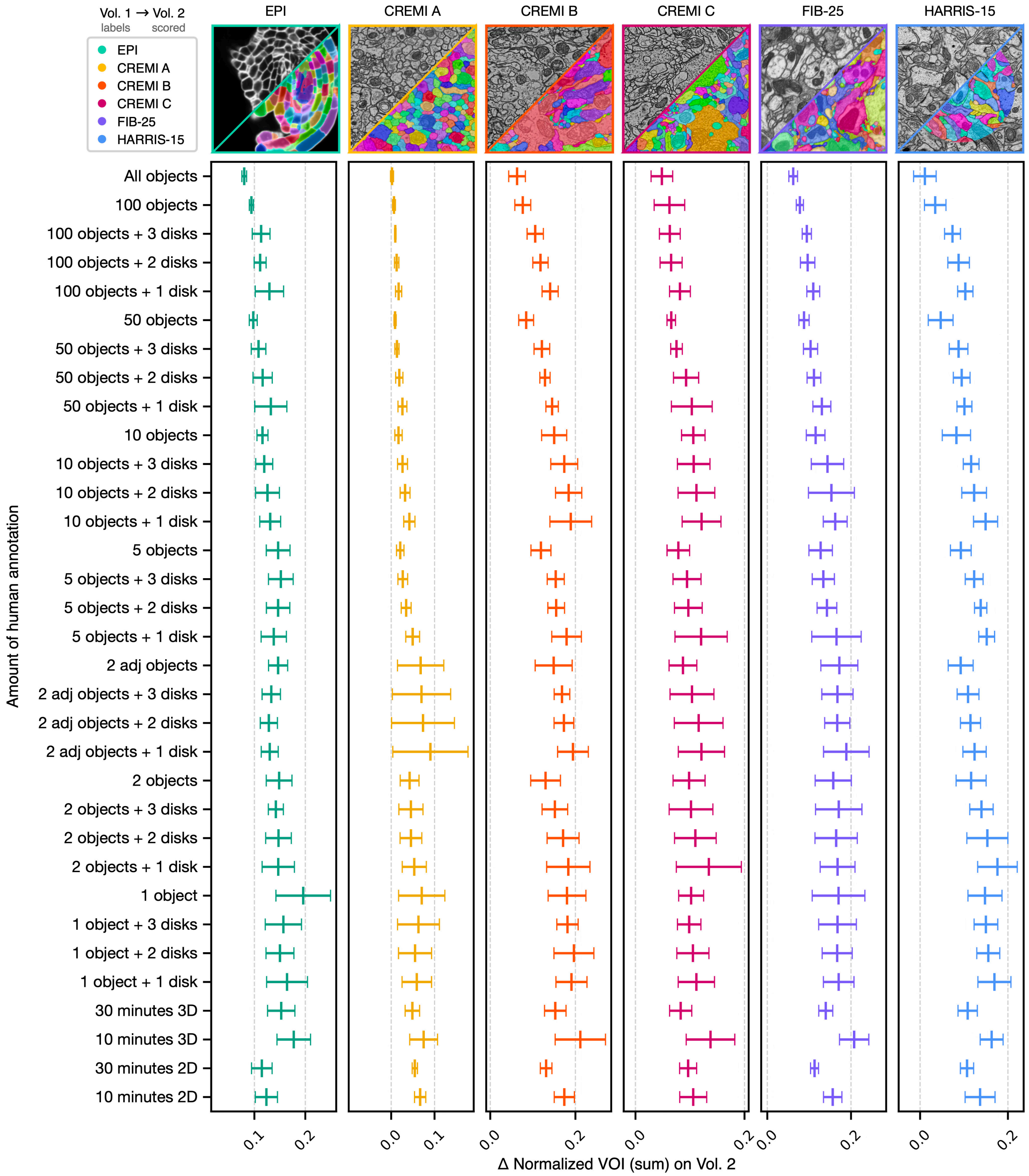


**Supplementary Fig. 3 | 2D→3D method trained on sparse labels of Volume 1 and scored on Volume 2: normalized VOI sum.** VOI, variation of information; LSD, local shape descriptor; MTLSD, multitask LSD. One panel per dataset (EPI, CREMI-A, CREMI-B, CREMI-C, FIB-25, HARRIS-15; the thumbnails show the raw image and the ground truth). Rows are the annotation levels used to train the 2D network: object-level sparsities with and without 1, 2 or 3 disks, and 10 or 30 minutes of non-expert 2D or 3D painting (Supplementary Note 1). Δ is the normalized VOI sum of the 2D→3D segmentation minus that of the 3D MTLSD network trained on the dense ground truth of the same repetition. Lower is better. Points are means and bars are ±1 s.d. over n = 12 differences (4 prediction types × 3 repetitions; Supplementary Note 4).

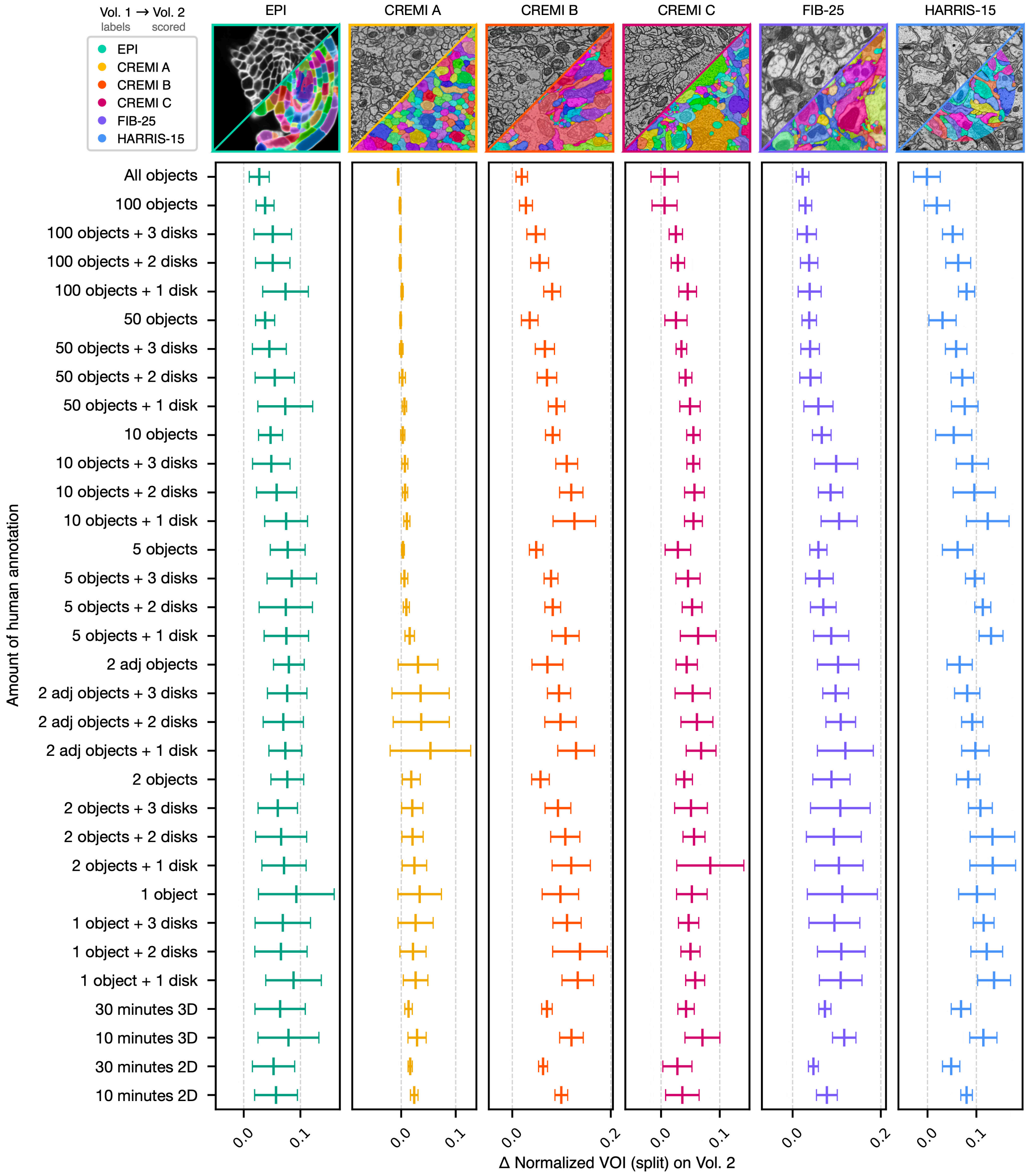


**Supplementary Fig. 4 | 2D→3D method trained on sparse labels of Volume 1 and scored on Volume 2: normalized VOI split.** VOI, variation of information; LSD, local shape descriptor; MTLSD, multitask LSD. One panel per dataset; rows are the annotation levels used to train the 2D network, as in Supplementary Fig. 3. Δ is the normalized VOI split of the 2D→3D segmentation minus that of the 3D MTLSD network trained on the dense ground truth of the same repetition. Lower is better. Points are means and bars are ±1 s.d. over n = 12 differences (4 prediction types × 3 repetitions; Supplementary Note 4).

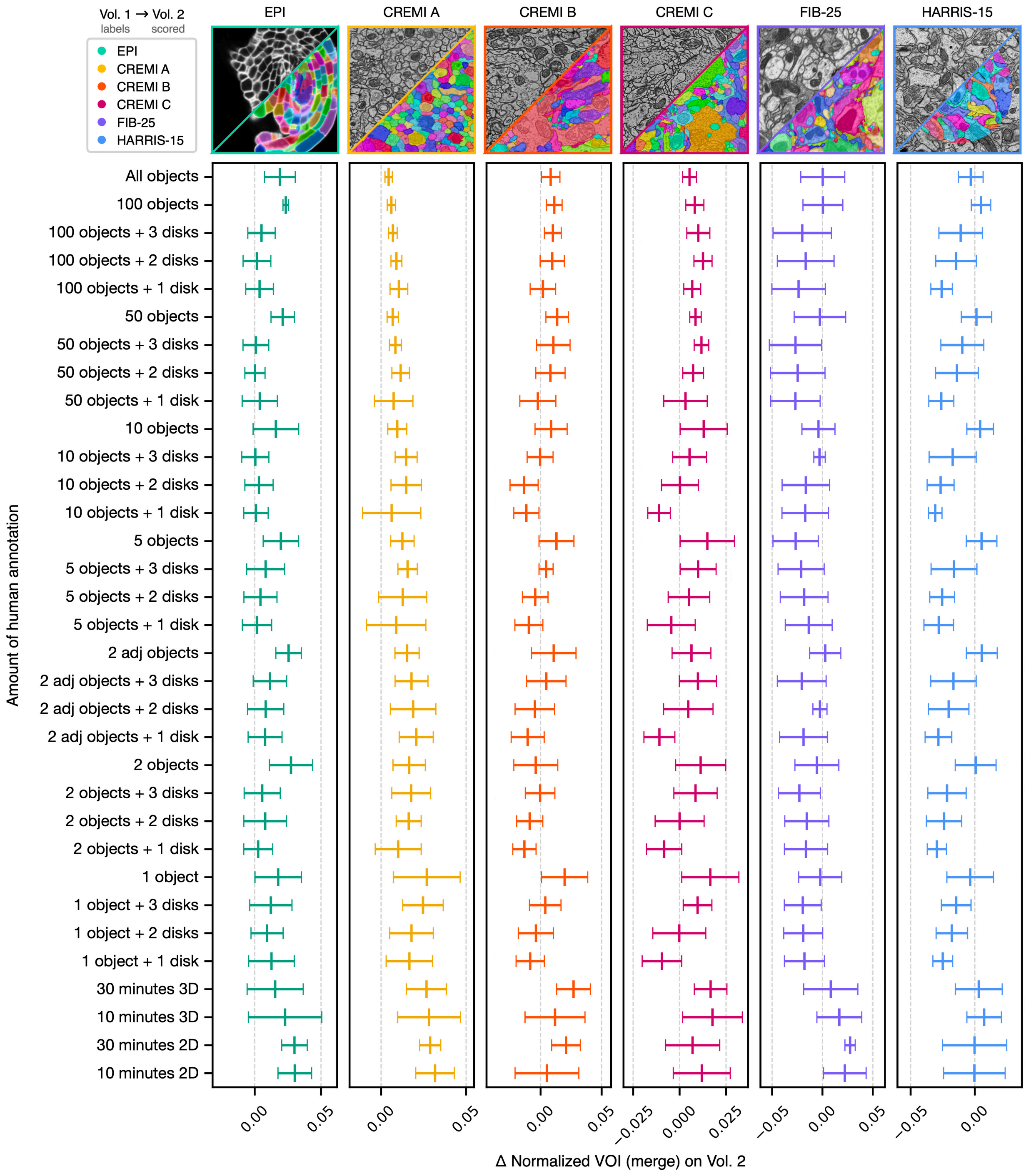


**Supplementary Fig. 5 | 2D→3D method trained on sparse labels of Volume 1 and scored on Volume 2: normalized VOI merge.** VOI, variation of information; LSD, local shape descriptor;

MTLSD, multitask LSD. One panel per dataset; rows are the annotation levels used to train the 2D network, as in Supplementary Fig. 3. Δ is the normalized VOI merge of the 2D→3D segmentation minus that of the 3D MTLSD network trained on the dense ground truth of the same repetition. Lower is better. Points are means and bars are ±1 s.d. over n = 12 differences (4 prediction types × 3 repetitions; Supplementary Note 4).

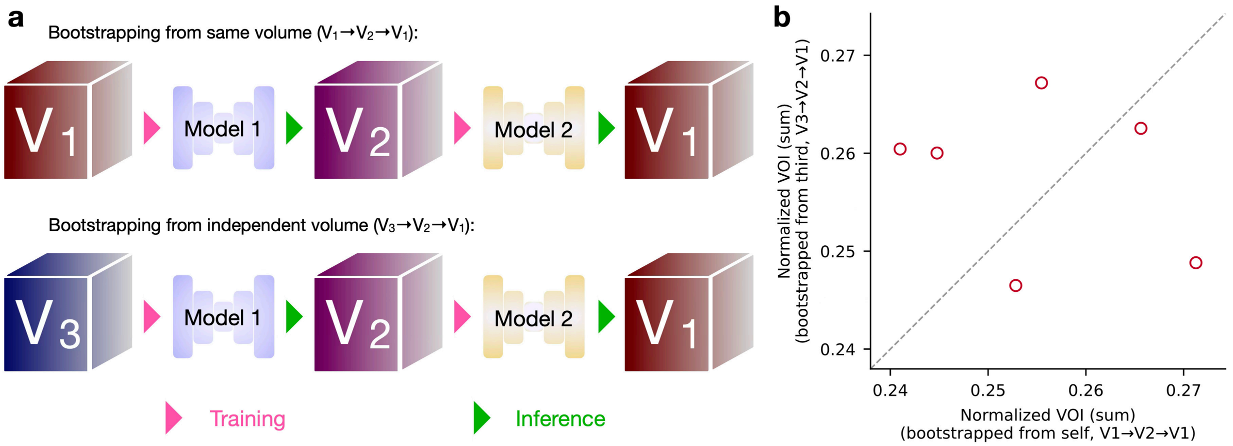


**Supplementary Fig. 6 | Comparison of bootstrapping paths on three z-slabs of HARRIS-15.** MTLSD, multitask LSD; VOI, variation of information. **a**, The two paths (Supplementary Note 5). Direct path (V1→V2→V1): a 3D MTLSD network trained on the dense labels of the evaluation slab V1 segments the intermediate slab V2. A second 3D MTLSD network trained on that unproofread segmentation segments V1. Control path (V3→V2→V1): the same chain, but the first network is trained on the dense labels of the independent slab V3. Pink arrowheads mark training and green arrowheads mark inference. **b**, Normalized VOI (sum) of the control path against that of the direct path for all six ordered pairs of the Apical slabs A, B and C (Supplementary Table 1). Each point is one pair (n = 6); there are no error bars. The dashed line marks equal scores.

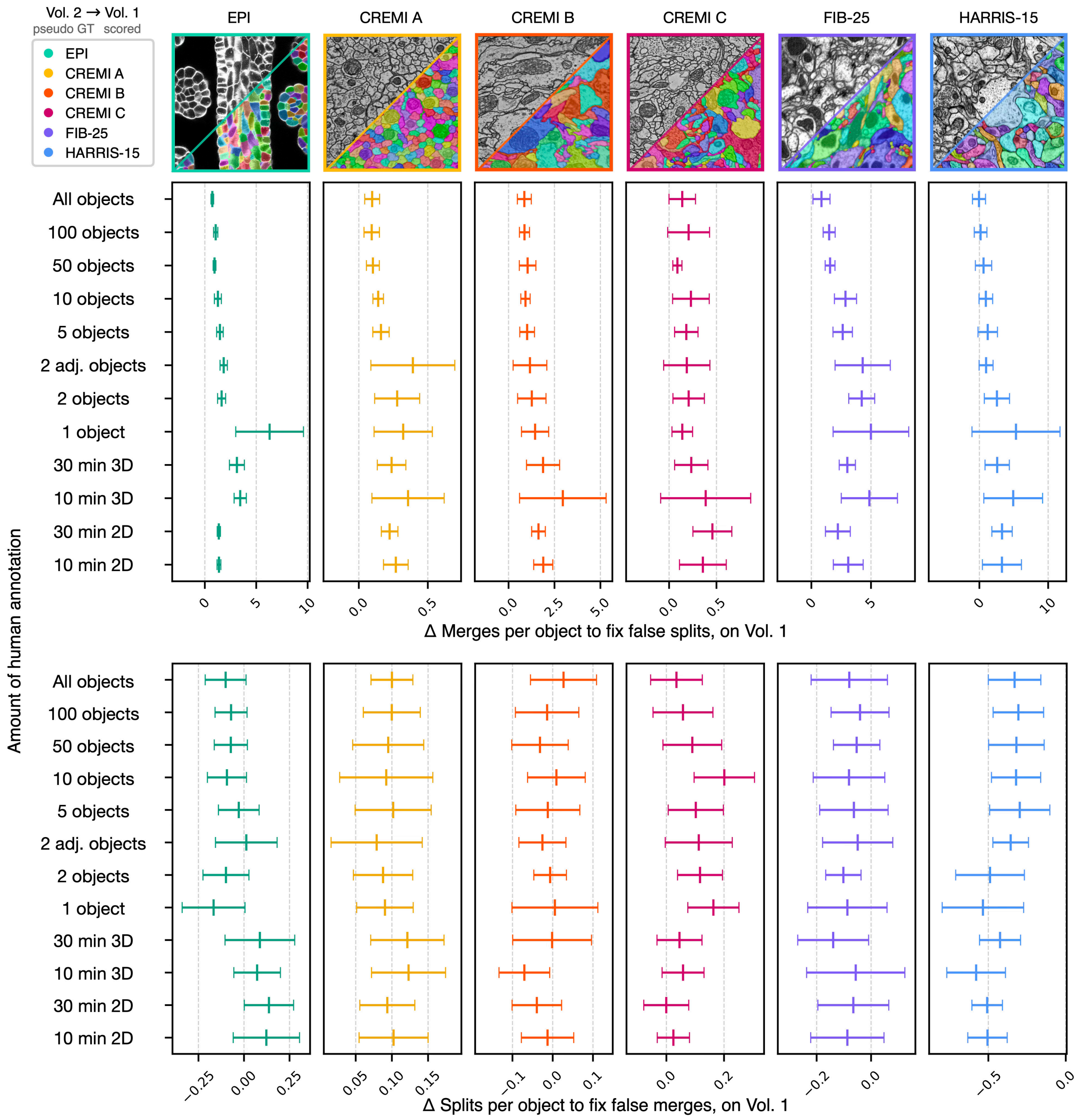


**Supplementary Fig. 7 | Bootstrapped 3D models trained on the 2D→3D pseudo ground truth of Volume 2 and scored on Volume 1: average edits per object needed to proofread.** MCM, min-cut metric; MTLSD, multitask LSD. One column of panels per dataset (EPI, CREMI-A, CREMI-B, CREMI-C, FIB-25, HARRIS-15). Rows are the annotation levels used to generate the pseudo ground truth for training the bootstrapped 3D MTLSD network (Supplementary Note 4). Top, Δ of the average number of merges per object needed to fix false splits; bottom, Δ of the average number of splits per object needed to fix false merges (MCM). Δ is the value of the bootstrapped network minus that of the non-bootstrapped 3D MTLSD network trained on the dense ground truth of the same repetition. Lower is better. Points are means and bars are ±1 s.d. over n = 12 bootstrapped models

(4 pseudo ground-truth network variants × 3 repetitions; one variant per prediction type of Supplementary Note 4).

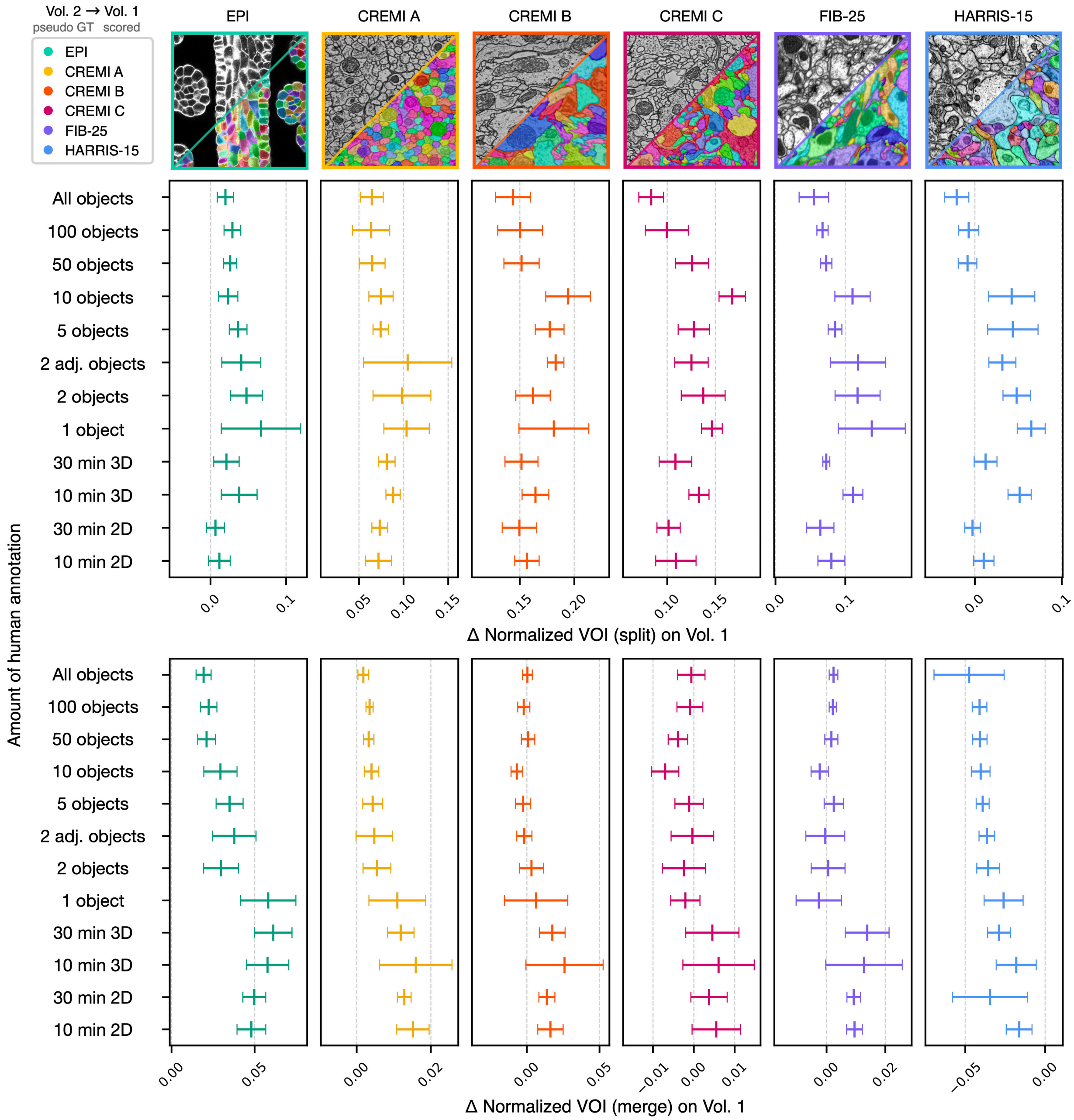


**Supplementary Fig. 8 | Bootstrapped 3D models trained on the 2D→3D pseudo ground truth of Volume 2 and scored on Volume 1: normalized VOI split and merge.** VOI, variation of information; MTLSD, multitask LSD. One column of panels per dataset; rows are the annotation levels used to generate the pseudo ground truth for training the bootstrapped 3D MTLSD network (Supplementary Note 4). Top, Δ normalized VOI split; bottom, Δ normalized VOI merge. Δ is the value of the bootstrapped network minus that of the non-bootstrapped 3D MTLSD network trained on the dense ground truth of the same repetition. Lower is better. The split panel is evaluated at the post-processing configuration with the fewest total edits, the merge panel at the configuration with the lowest VOI sum. Points are means and bars are ±1 s.d. over n = 12 bootstrapped models (4 pseudo ground-truth network variants × 3 repetitions; one variant per prediction type of Supplementary Note 4).

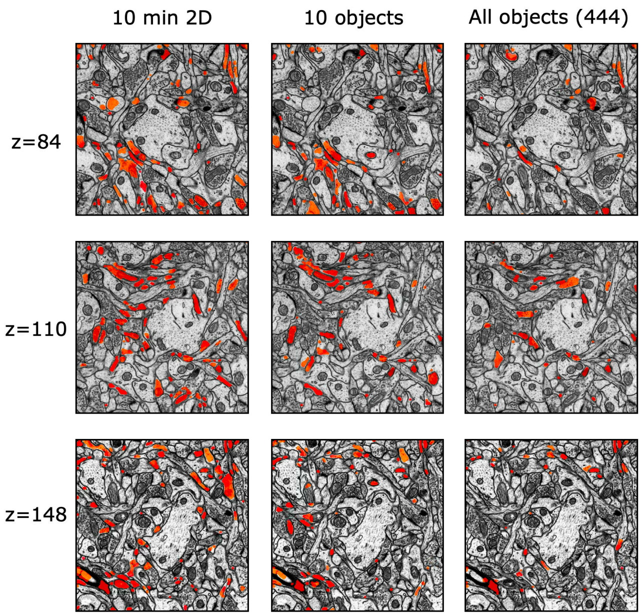


**Supplementary Fig. 9 | LSD inconsistencies of bootstrapped HARRIS-15 segmentations.** LSD, local shape descriptor; MTLSD, multitask LSD. LSD inconsistencies (red) are element-wise differences between LSDs predicted by the bootstrapped 3D MTLSD network and those computed from its final segmentation. These differences mark regions where residual errors are likely. Columns, bootstrapped networks trained on pseudo ground truth from 10 minutes of 2D annotation, 10 objects and all objects (444). Rows, three sections of HARRIS-15. Illustration only; no quantification.

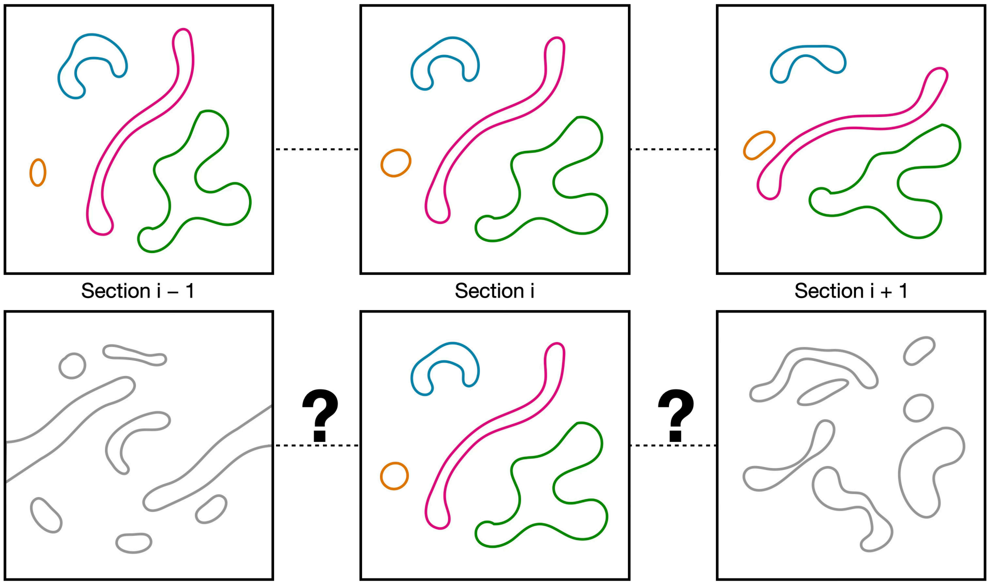


**Supplementary Fig. 10 | Insufficient z resolution results in ambiguous inter-section connectivity.** Three consecutive sections (i − 1, i, i + 1) of mock image data. Top, sufficient resolution between sections: the colored outlines can be followed from section to section. Bottom, insufficient resolution between sections (gray outlines, question marks): even human annotators can struggle to determine which segments connect across sections. Schematic; no data.
